# Prefusion-specific glycoprotein B human antibodies protect against neonatal HSV-2 infection

**DOI:** 10.64898/2026.02.23.707514

**Authors:** Parastoo Amlashi, Jeongryeol Kim, Nisali N. Mendis, Perry T. Wasdin, Lindsay E. Bass, Gwen Jordaan, Alexandra A. Abu-Shmais, Matthew D. Slein, Nicole V. Johnson, Rachel H. Bonami, David A. Leib, Margaret E. Ackerman, Jason S. McLellan, Ivelin S. Georgiev

**Author notes:** These authors contributed equally to this work (Parastoo Amlashi, Jeongryeol Kim).

## Abstract

Herpes simplex virus (HSV) causes severe neonatal infections and long-term neurological issues, with antivirals offering limited efficacy and no approved vaccines. Viral entry requires glycoprotein B (gB) for membrane fusion, yet no human antibodies to the prefusion form of gB have been identified, and antibodies targeting vulnerable epitopes are limited. We used prefusion-stabilized HSV-2 gB and LIBRA-seq technology to analyze B cells from seropositive and healthy donors, discovering four prefusion-specific human antibodies binding unique gB epitopes as determined by cryo-EM. Antibodies 5-18 and 3-6 showed strong cross-neutralization against HSV-1 and HSV-2, offering protection in a neonatal mouse model. These findings highlight the molecular and structural determinants of HSV-2 prefusion gB recognition by human antibodies and demonstrate the translational potential of prefusion-specific gB antibodies.

## INTRODUCTION

Herpes simplex viruses (HSV) are enveloped, double-stranded DNA alphaherpesviruses that establish lifelong infection in the peripheral nervous system and cause mucocutaneous disease (*1, 2*). HSV-1 typically presents as orolabial infection, whereas HSV-2 is associated with recurrent genital lesions and episodic viral shedding (*3*). Globally, an estimated 13% of individuals aged 15–49 are infected with HSV-2, highlighting its persistent circulation and high seroprevalence (*4*). Beyond mucocutaneous disease, HSV-2 contributes significantly to morbidity through neurologic complications and severe outcomes in neonates and immunocompromised individuals (*5*). Neonatal HSV-2 infection, most commonly resulting from vertical transmission during birth, can cause long-term neurological impairment or death (*6*). Current treatment relies on nucleoside analogs such as acyclovir, and although high-dose regimens improve survival in neonates, the risks of neurodevelopmental sequelae and relapse remain considerable (*7, 8*). Despite antiviral therapy, disseminated neonatal HSV-2 infection retains a mortality rate of approximately 30% (*7, 9*), underscoring the urgent need for both preventive and adjunctive therapies. In this context, monoclonal antibodies (mAbs) targeting essential viral glycoproteins represent a promising avenue for therapeutic development.

HSV entry depends on a coordinated mechanism involving the surface glycoproteins gB, gD, and the gH/gL complex (*10*). Entry is initiated by the receptor-binding gD, which activates the gH/gL heterodimer to trigger the membrane fusogen gB, a homotrimer that drives fusion of the viral and host cell membranes (*11–13*). gB is highly conserved across herpesviruses and plays a critical role in viral attachment and fusion, making it an attractive target for the development of vaccines and antibodies (*3, 14*). As a class III fusion protein, gB undergoes a large-scale conformational rearrangement from a metastable prefusion state to a stable postfusion state (*15, 16*). Stabilizing gB in its prefusion state provides a promising strategy for eliciting potent neutralizing antibody responses, as the prefusion conformation presents unique antibody-accessible sites that are concealed or disrupted in the postfusion conformation, as demonstrated for other viral fusion proteins (*17–23*). Indeed, antibodies targeting prefusion-specific epitopes on other viral fusion proteins have been shown to exhibit potent neutralizing activity (*19–23*). However, the structural transition of gB primarily involves domain rearrangements rather than the secondary-structure refolding characteristic of class I fusion proteins, and stabilizing the prefusion state has proven technically challenging, hindering the isolation of prefusion-specific gB antibodies (*24, 25*). Although a camelid nanobody (*26*) and a murine mAb (*25*), both elicited by immunization with recombinant antigens, were recently shown to specifically recognize HSV prefusion gB, it remains unknown whether natural human infection elicits similar prefusion-specific antibody responses and whether such antibodies confer potent neutralization or protection from morbidity and mortality.

Although several mAbs protect against HSV-1/2 in preclinical models, most target gD; only a few target gB, fewer still are human-derived, and none are prefusion-specific (*27–32*). The gB ectodomain consists of five structural domains, and most potently neutralizing gB-specific antibodies recognize domains I, II, and IV (**fig. S1**) (*33–35*). Notably, HDIT101, a humanized mAb targeting domain I, did not demonstrate superiority over standard-of-care therapy in a phase 2 clinical trial for recurrent genital herpes (*36, 37*). These findings underscore the importance of further delineating the antigenic landscape of prefusion gB and comprehensively profiling the human B-cell response to HSV-2, both of which remain critical for informing next-generation vaccine development and antibody-based interventions.

To address these knowledge gaps, we characterized the HSV-2 gB-specific B-cell repertoire in HSV-2 seropositive individuals and healthy individuals with no recorded history of HSV infection. Using the LIBRA-seq (Linking B cell Receptor Sequence to Antigen Specificity through Sequencing) antibody-discovery technology (*38*), we identified four prefusion-specific and two prefusion-preferring antibodies from human peripheral blood mononuclear cell (PBMC) samples. Two prefusion-specific antibodies, 5-18 and 3-6, exhibited potent cross-neutralizing activity against clinical isolates of both HSV-1 and HSV-2. To define the structural basis of their specificity, we determined cryo-electron microscopy (cryo-EM) structures of a prefusion-stabilized HSV-2 gB ectodomain in complex with the antigen-binding fragments (Fabs) of each antibody. Fab 5-18, along with another prefusion-specific Fab 1-14, recognize quaternary epitopes that span domains I and IV of adjacent protomers, whereas Fab 3-6 binds a distinct epitope that bridges domains I and II and the intervening hinge region. Finally, both 5-18 and 3-6 conferred protection in a well-established neonatal mouse model of HSV-2 infection, underscoring their potential as candidate antibody-based interventions to prevent neonatal HSV-2 infection and associated mortality.

## RESULTS

### Identification of a diverse set of HSV-2 prefusion gB-specific mAbs by LIBRA-seq

We applied LIBRA-seq to identify HSV-2 prefusion gB-specific B cells from human donor PBMCs. The LIBRA-seq antigen panel included stabilized prefusion and postfusion gB constructs from several clinically relevant human herpesviruses, including HSV-2, human cytomegalovirus (HCMV), and Epstein-Barr virus (EBV), as well as control antigens HIV BG505 gp140 V9.3 SOSIP and influenza HA NC99, intended to filter out cells with non-specific binding activity. Prefusion-stabilized gB constructs used were HSV-2 gB-G3 (*39*), HCMV gB C7 (*40*) (PDB ID: 7KDP), EBV gB C3-GT (*41*) (PDB ID: 9OAL), as well as the postfusion gB ectodomain base constructs of each herpesvirus. After LIBRA-seq sequencing and computational filtering as previously described (*38*), we isolated a total of 449 antigen-specific B cells from eight donors: three with confirmed HSV-2 seropositivity, three with confirmed HCMV seropositivity, and two healthy donors with no recorded history of HSV or HCMV. A LIBRA-seq score (LSS) of 1 or greater was selected as a cutoff to identify B cells predicted to have specificity for HSV-2 gB. A total of 69 B cells had an LSS ≥1 for HSV-2 prefusion gB, and 75 had an LSS ≥1 for HSV-2 postfusion gB, with some B cells having an LSS of ≥1 for both. Of these, we initially selected 22 clonally distinct antibodies with diverse sequence characteristics and varying LIBRA-seq scores for HSV-2 prefusion gB and postfusion gB (**Fig. 1A**).

**Fig 1.**
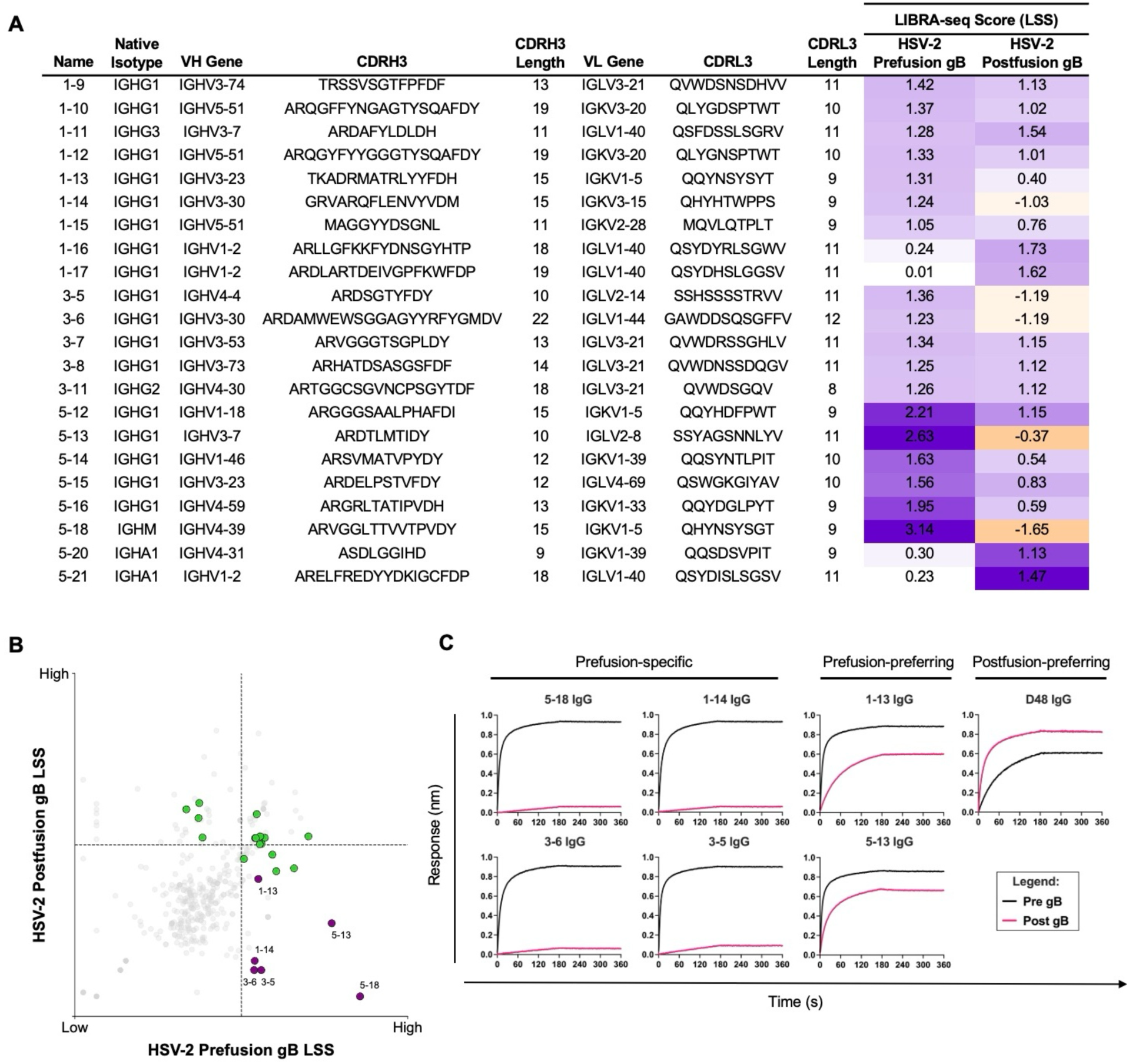
Identification and characterization of HSV-2 prefusion-specific human antibodies. **(A)** Sequence properties of HSV-2 gB antibodies identified by LIBRA-seq. The complementarity-determining region (CDR) sequences and lengths for each antibody are shown at the amino acid level. LIBRA-seq scores (LSS) for the selected antibodies to HSV-2 pre- and postfusion gB antigens are shown as a heat map from –2 (tan) to 0 (white) to 2 (purple); scores outside of this range are shown as the minimum and maximum values. **(B)** LIBRA-seq scores for each recovered B cell are plotted, with cutoffs set at an LSS of 1. Antibodies encoded by B cells colored in purple were selected for experimental validation. The remaining 22 selected antibodies are colored green. **(C)** BLI sensorgrams for immobilized IgGs binding to prefusion-stabilized HSV-2 gB (black) or postfusion HSV-2 gB (pink).

These 22 antibodies were produced recombinantly as IgG1 mAbs and screened for binding by indirect enzyme-linked immunosorbent assay (ELISA). The selected antibodies bound either prefusion gB, postfusion gB, or both (**fig. S2A**). To further explore the potential for identifying prefusion-specific human antibodies against HSV-2 gB, we specifically focused on six antibodies that showed: (1) high prefusion and low postfusion gB LIBRA-seq scores (**Fig. 1B**) and/or (2) stronger antibody binding for prefusion than postfusion gB by ELISA (**fig. S2B**). Biolayer interferometry (BLI) experiments were performed to assess the conformational specificity of the isolated antibodies. In these experiments, the IgGs were immobilized to anti-human IgG Fc Capture tips and assessed for binding to prefusion- and postfusion-stabilized HSV-2 gB ectodomains at 200 nM, a concentration sufficient to observe high-affinity interactions. Antibodies 5-18, 1-14, 3-6, and 3-5 each displayed saturated binding to prefusion-stabilized gB but negligible binding to postfusion gB, indicative of prefusion-specific binding (**Fig. 1C**). Antibodies 1-13 and 5-13 bound both proteins well, but with a faster on-rate for prefusion gB, indicative of a preference for the prefusion conformation. Conversely, anti-HSV gB antibody D48 (*42*) bound both pre- and postfusion gB but had a faster on-rate for postfusion gB. Overall, four antibodies were classified as prefusion-specific and two as prefusion-preferring based on their reactivity to the two stabilized ectodomains, with the BLI data generally in agreement with LIBRA-seq results.

To assess how common the HSV-2 gB prefusion-specific and prefusion-preferring antibodies are in the general population, these mAb sequences were compared against a general database of 1.7 million paired antibodies (**fig. S3A**). Several prefusion-specific mAbs exhibited matches with ≥70% HCDR3 identity, most prominently mAb 3-5 (4,498 matches), with fewer matches observed for 5-18 (20 matches) and 3-6 (2 matches), and none for 1-14 (**fig. S3B**). However, no prefusion-specific mAbs shared both heavy and light chain V gene usage with antibodies in the database. Although mAb 5-18 showed multiple VH-matched pairs with high HCDR3 identity, none met the criteria for full clones due to a lack of shared VL usage. Among prefusion-preferring mAbs, 5-13 yielded 163 matches and included a single full clone with matched VH and VL genes and ≥70% CDR3 identity, whereas mAb 1-14 showed no such matches (**fig. S3B**). Together, these analyses suggest that no true public clonotypes could be identified for the prefusion-specific gB mAbs from our study.

### Prefusion-specific gB antibodies potently neutralize HSV

To investigate epitope groups on prefusion HSV-2 gB, we performed a pairwise competition ELISA with all 22 antibodies initially identified from the dataset (**Fig. 2A**). The patterns of binding-inhibition among antibody pairs compared to signal with no antibody competition suggested the existence of at least one shared epitope on prefusion gB, as well as clusters of antibodies that target overlapping or closely positioned binding regions (**Fig. 2B**). To further characterize these interactions, we measured the binding affinities of the Fabs to prefusion-stabilized HSV-2 gB via BLI (**Fig. 2C and fig. S4**). Both prefusion-specific and prefusion-preferring mAbs exhibited picomolar affinities for prefusion-stabilized HSV-2 gB. Among them, Fab 1-14 showed the strongest binding, with an equilibrium dissociation constant (*K*_d_) of 0.03 nM and a dissociation rate constant (*k*_off_) at the instrument’s limit of detection (1.2 × 10⁻^5^ s⁻^1^). Prefusion-specific Fab 3-6 exhibited a *K*_d_ of 0.12 nM and a *k*_off_ of 5.3 × 10⁻^5^ s⁻^1^. Given the high sequence similarity between HSV-2 and HSV-1 gB, we produced a prefusion-stabilized HSV-1 gB ectodomain construct by applying the same stabilizing substitutions developed for HSV-2 gB (**fig. S5 and fig. S6A-C**). Binding affinities of the antibodies for the prefusion-stabilized HSV-1 gB were then determined via BLI by immobilizing HSV-1 prefusion gB to Ni^2+^-NTA tips and dipping into Fab (**Fig. 2C and fig. S6D**). Fab 3-6 exhibited a slightly improved affinity (*K*_d_ = 0.08 nM), driven by a faster association rate constant (*k*_on_ = 9.8 × 10⁵ M⁻¹s⁻¹). In contrast, Fab 5-18 showed substantially reduced binding to HSV-1 gB, with a faster dissociation rate constant and lower overall affinity (*K*_d_ = 40 nM). Fabs 1-14 and 3-5 bound HSV-1 gB with affinities comparable to those observed for HSV-2 gB.

**Fig 2.**
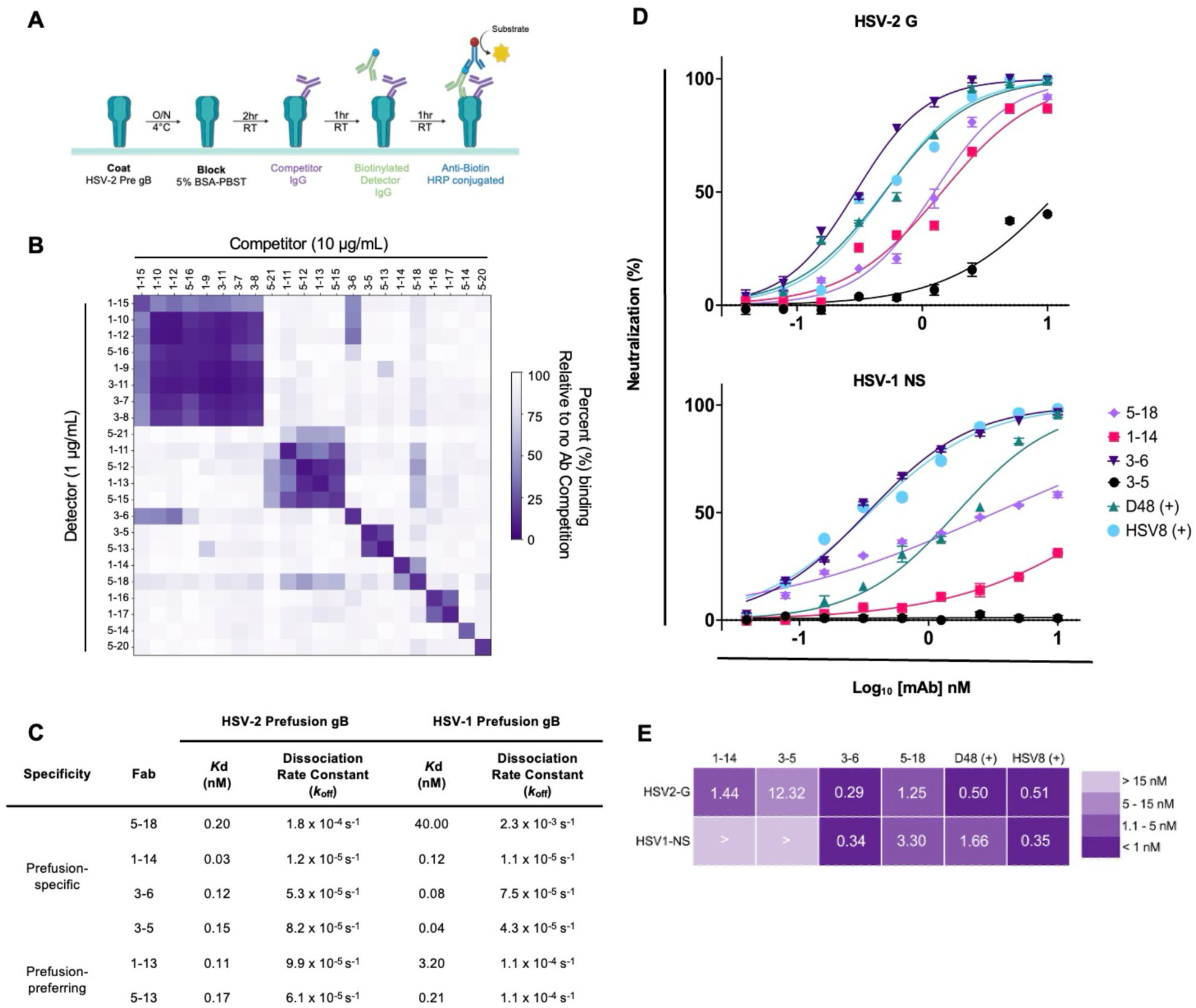
Binding and neutralization of prefusion-specific gB antibodies. **(A)** ELISA-based antibody-antibody binding competition assay (schematic created using Biorender.com). **(B)** The percentage of binding remaining in the presence of a competitor mAb is shown for each mAb, with darker colors corresponding to stronger competition between the given pair of mAbs. Dark purple clusters indicate a group of mAbs targeting overlapping epitopes on the HSV-2 prefusion gB antigen. **(C)** Equilibrium dissociation constants (Kd) and dissociation rate constants (kd) of gB Fabs binding to immobilized HSV-2 prefusion gB on Ni2+-NTA biosensors. **(D)** Neutralization profiles of gB-specific mAbs against HSV-2 G and HSV-1 NS strains detected by a plaque reduction assay. **(E)** Neutralization midpoint concentration (IC50) summary heatmap, with strong neutralization of HSV-2 G and HSV-1 NS (< 1 nM) shown in dark purple and weak/non-neutralizing (> 15 nM) shown in light purple. Calculated by non-linear regression analysis using GraphPad Prism software.

We next assessed the neutralization activity of the mAbs against the prototypic HSV-2 strain G via plaque-reduction assay (**Fig. 2D**). mAb 3-6 exhibited the most potent neutralization (IC_50_ = 0.29 nM), exceeding well-characterized antibodies such as D48 (IC_50_ = 0.50 nM) and HSV8 (an anti-gD antibody; IC_50_ = 0.51 nM). mAb 5-18 (IC_50_ = 1.25 nM) and 1-14 (IC_50_ = 1.44 nM) were also potent and showed comparable neutralizing activity (**Fig. 2E**). In contrast, 3-5 was relatively weakly neutralizing despite having an affinity similar to the other prefusion-specific antibodies. We also evaluated the neutralization activity of these prefusion-specific antibodies against the clinical HSV-1 strain NS. Consistent with results for HSV-2, mAb 3-6 exhibited the most potent neutralizing activity (IC_50_ = 0.34 nM), exceeding that of D48 (IC_50_ = 1.66 nM). mAb 5-18 (IC_50_ = 3.30 nM) was the next most potent, followed by 1-14 (IC_50_ = 33.46 nM). In contrast, 3-5 did not neutralize HSV-1 NS infection at the concentrations tested. These findings demonstrate that HSV-2 prefusion-specific gB antibodies can cross-react with HSV-1 gB, and that 3-6 is the most potent antibody against both HSV-1 and HSV-2.

We next evaluated *in vitro* autoreactivity of the four prefusion-specific gB antibodies to self-antigens using immortalized human epithelial cell lines. We determined intracellular and extracellular IgG binding to live, permeabilized HEp-2 and HEK293T cells by flow cytometry. At 25 µg/mL, none of the prefusion-specific gB antibodies displayed extracellular binding to either cell line (**Table S1**). Although mAbs 1-14 and 5-18 exhibited low levels of intracellular binding compared to the positive control antibodies 4E10 and 2F5, no such reactivity was observed for mAbs 3-5 and 3-6 (**fig. S7**).

### Antibodies 5-18 and 1-14 recognize quaternary epitopes spanning domains I and IV of adjacent protomers

We determined cryo-EM structures of prefusion-stabilized HSV-2 gB ectodomain in complex with Fabs 5-18 and 1-14 at 2.9 Å and 3.2 Å resolution, respectively, to investigate the molecular basis of their prefusion-specific epitopes (**Fig. 3**, **fig. S8 and fig. S9**). Both antibodies recognize quaternary epitopes spanning domains I and IV of adjacent protomers, but through distinct interaction modes. Fab 5-18 makes extensive contacts with both domains, burying a total surface area of 1,440 Å^2^, with 835 Å^2^ on domain I and 605 Å^2^ on domain IV (**Fig. 3A-B and fig. S10A**). The antibody forms 14 hydrogen bonds and 4 salt bridges with domain I and 10 hydrogen bonds with domain IV (**Table S2**). All three complementarity-determining regions of the heavy chain (HCDRs) engage domain I. HCDR1 and HCDR2 recognize domain I, whereas HCDR3 spans domains I and IV (**fig. S10B**). Complementarity-determining regions of the light chain (LCDRs) 1 and 3 primarily interact with domain I, whereas LCDR2 interacts extensively with domain IV, establishing a continuous hydrogen-bond network that stabilizes the interface (**fig. S10C).** This distribution of the epitope on to adjacent protomers suggests that binding of 5-18, as well as 1-14, prevents protomer separation and locks gB in the prefusion conformation, providing a mechanistic basis for neutralization. Mapping the epitope of 5-18 on to the structure of postfusion gB (PDB ID: 8RH2) reveals that the epitope is disrupted in the postfusion conformation due to the reorientation of domain I relative to domain IV, resulting in contact residues on gB separated by approximately 100 Å (**Fig. 3C**). Additionally, some of the epitope residues on domain I are buried by domain V of an adjacent protomer in the postfusion conformation, collectively providing a structural basis for the prefusion specificity of 5-18.

**Fig 3.**
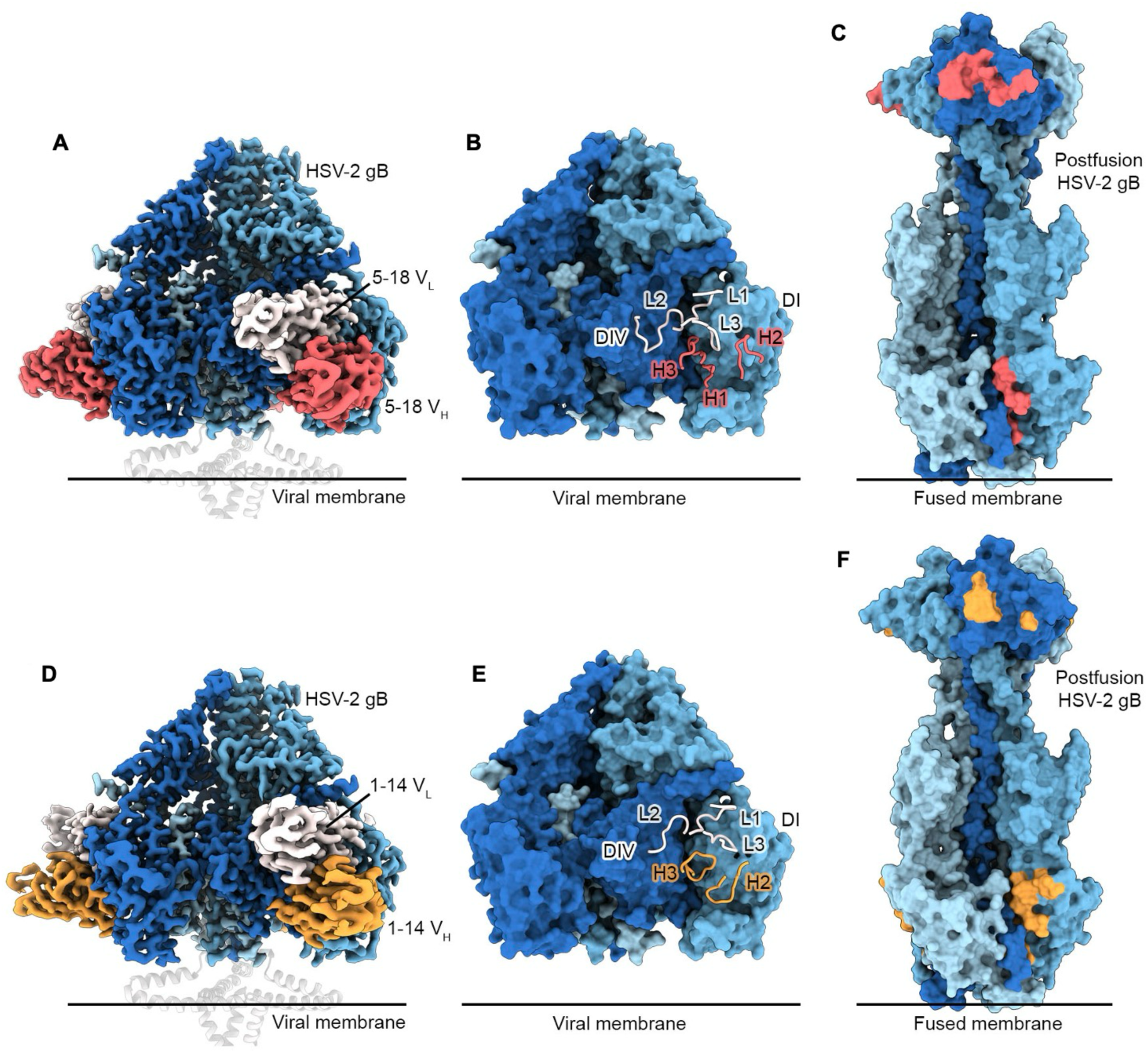
Fabs 5-18 and 1-14 recognize prefusion-specific quaternary epitopes spanning domains I and IV. **(A)** Cryo-EM map side view of Fab 5-18 bound to the HSV-2 prefusion gB trimer. The 5-18 VH is colored red, and the VL is colored white. Each prefusion gB protomer is colored a different shade of blue. The MPR and TM regions from PDB ID: 9Q9L are shown as transparent helices, and a horizontal black line indicates the approximate position of the viral membrane. **(B)** Side view of prefusion HSV-2 gB depicted as a molecular surface with the 5-18 CDRs depicted as ribbons. Domain I (DI) and domain IV (DIV) are labeled on gB. **(C)** Side view of postfusion HSV-2 gB (PDB ID: 8RH2) shown as a molecular surface with the 5-18 epitope colored red. The fused membrane is indicated by a horizontal black line. **D, E, and F,** are the same as **A, B,** and **C,** but depict the Fab 1-14 complex, CDRs, and epitope.

Fab 1-14 has a smaller binding footprint than 5-18, burying a total surface area of 1,125 Å^2^ (**Fig. 3D-E** and **fig. S10D**). Notably, interactions with domain IV differ substantially: 1-14 buries 244 Å^2^ on domain IV versus 605 Å^2^ for 5-18. The majority of the 1-14 contacts are mediated by HCDR2 and HCDR3, which make extensive interactions with the long loop between β14 and β15 in domain I (**fig. S10E**). LCDR1 and LCDR3 primarily contact domain I, whereas LCDR2 and framework 3 (FR3) make limited contacts with domain IV (buried surface area of 121 Å^2^), including residues in the N-terminal α1 helix (**fig. S10F**). Similar to 5-18, the epitope of 1-14 becomes discontinuous and partially buried in the postfusion conformation, providing the structural basis for its prefusion-specificity (**Fig. 3F**).

Interestingly, the epitopes of 5-18 and 1-14 closely resemble the epitope targeted by a recently described prefusion-specific gB-directed camelid nanobody Nb1_gbHSV (*26*), which recognizes a quaternary epitope spanning domains I, III, and IV (**fig. S11**). An epitope comparison shows that the Nb1_gbHSV contacts many of the same residues as Fabs 5-18 and 1-14, but the nanobody reaches more deeply into the space between domains I and IV, contacting domain III (**fig. S11E-F**). By contrast, Fabs 5-18 and 1-14 form more extensive contacts across domains I and IV (**fig. S11A-D**). These structural studies reveal multiple solutions by the human and camelid immune systems to recognize a similar quaternary epitope on prefusion gB proteins.

### Antibodies 3-6 and 3-5 bind distinct epitopes located within a single protomer

Next, we determined a cryo-EM structure of a ternary complex of prefusion-stabilized HSV-2 gB bound to Fabs 3-6 and 3-5 at a resolution of 3.1 Å (**Fig. 4A and fig. S12**). Fab 3-6 engages an extensive interdomain epitope encompassing domains I, II, and the hinge region connecting them, burying a total surface area of 886 Å² (**Fig. 4B and fig. S13A**). Binding is mediated primarily by the heavy chain (806 Å²), with a much smaller contribution from the light chain (80 Å²), involving a total of 16 hydrogen bonds and 1 salt bridge (**Table S3**). The FR1 and HCDR1 of the heavy chain interact with the α4 helix of domain II. HCDR1, HCDR2, and HCDR3 form multiple hydrogen bonds with the backbone of hinge residues 351–358, establishing an extensive interaction network (**fig. S13B**). Additionally, Trp100_HCDR3_ is inserted into a hydrophobic pocket in domain I formed by Tyr152, Tyr279, Phe295, and Val352, burying 134 Å² of surface area (**Fig 4B and fig. S13C**).

**Fig 4.**
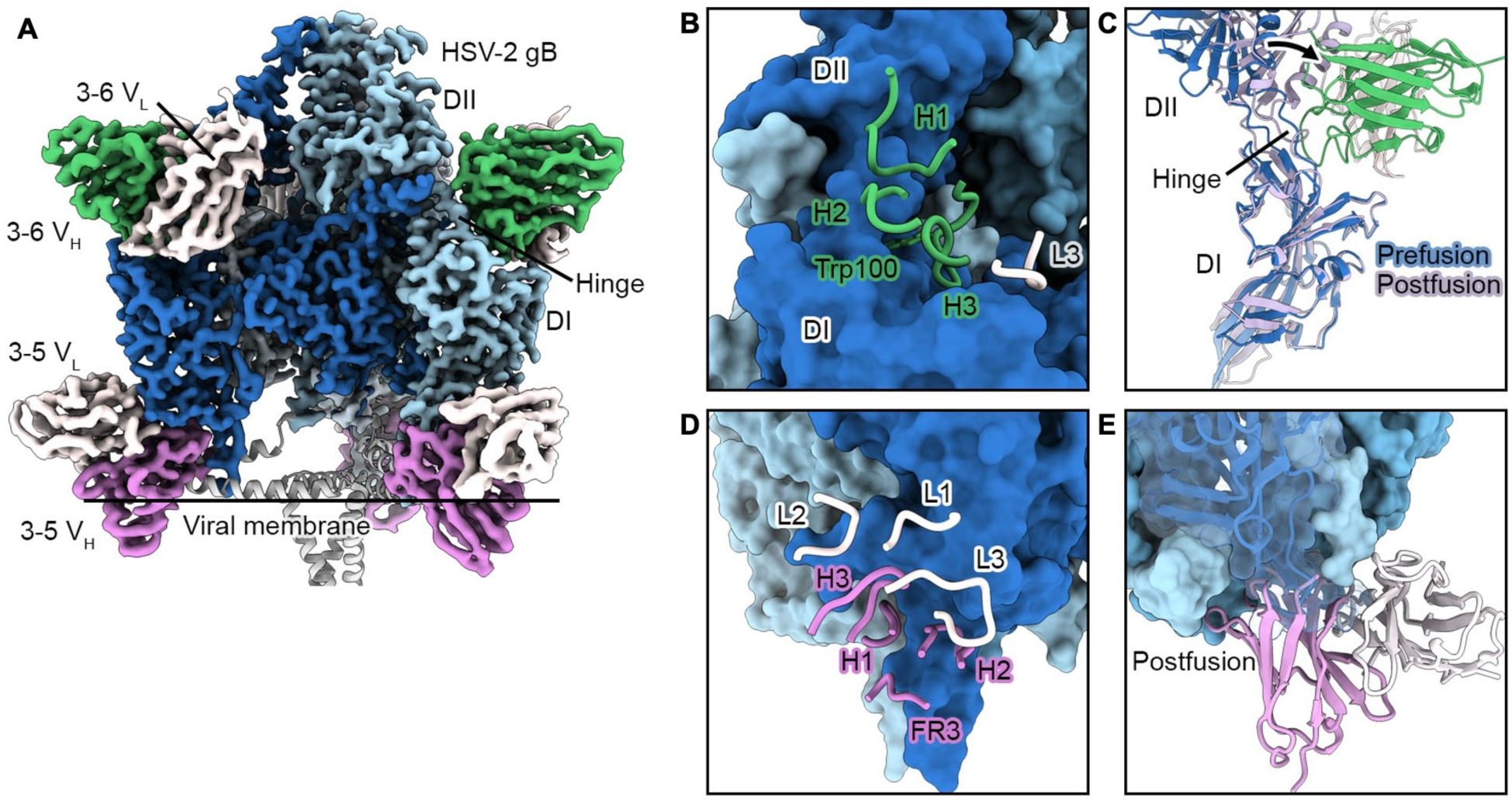
Antibodies 3-6 and 3-5 bind distinct prefusion-specific epitopes located within a single protomer. **(A)** Cryo-EM map side view of Fabs 3-6 and 3-5 bound to the HSV-2 prefusion gB trimer. The 3-6 VH is colored green, the 3-5 VH is colored magenta, and both VLs are colored white. Each prefusion gB protomer is colored a different shade of blue. The MPR and TM regions from PDB ID: 9Q9L are shown as white helices, and a horizontal black line indicates the approximate position of the viral membrane. **(B)** Magnified view of prefusion HSV-2 gB depicted as a molecular surface with the 3-6 CDRs depicted as ribbons. Domain I (DI) and domain II (DII) are labeled on gB. The sidechain of Trp100 in the HCDR3 (H3) is shown as sticks. **(C)** Superimposition of DI from the 3-6-bound prefusion gB monomer and a monomer from postfusion gB (PDB ID: 8RH2) showing the rearrangement of DII (indicated with a black arrow) that blocks Fab 3-6 binding. **(D)** Magnified view of prefusion HSV-2 gB depicted as a molecular surface with the 3-5 CDRs and framework 3 (FR3) region depicted as ribbons. **(E)** Magnified view of a superposition of the postfusion gB trimer (PDB ID: 8RH2) and Fab 3-5, with one protomer surface rendered transparent and the chain shown as ribbons. In the postfusion conformation, gB cannot bind Fab 3-5 due to numerous clashes.

Superposition of domain I from a Fab 3-6-bound protomer onto domain I of the postfusion gB structure shows that domain II reorients in the postfusion conformation and would clash with Fab 3-6 (**Fig. 4C**). The flexible hinge residues at the center of the epitope also undergo a conformation change, distorting the epitope. Collectively, these rearrangements provide a structural basis for the inability of 3-6 to bind trimeric postfusion gB. A recently reported prefusion gB-specific murine mAb named WS.HSV-1.24 (*25*) was shown to recognize a highly similar epitope, but it is rotated about the heavy chain such that the light chain is on the opposite side relative to that of 3-6 (**fig. S14**). Overall, 3-6 targets a highly neutralization-sensitive prefusion-specific epitope encompassing the hinge region between domains I and II, establishing this site as a vulnerable region on HSV-2 gB.

Antibody 3-5 binds to an epitope at the tip of domain I, a region not previously known to be targeted by gB-directed antibodies (**Fig. 4A**). This interaction enabled modeling of the domain I residues near the fusion loops, which were disordered in the HSV-2 gB structures complexed with 5-18 or 1-14. Fab 3-5 forms a large interface on domain I, burying a total of 1,296 Å^2^ of surface area. The heavy chain buries 980 Å^2^, whereas the light chain buries only 316 Å^2^. All heavy chain CDRs, as well as FR1 and FR3, participate in the interaction with domain I, forming 16 hydrogen bonds (**Table S3**). Residues in LCDR1 and LCDR3 form 7 hydrogen bonds with residues in domain I, further extending the interaction surface (**Table S3**). Interestingly, the angle of approach of Fab 3-5 is oriented upward from the membrane-proximal side, likely preventing binding to the native, membrane-embedded prefusion gB trimer on virions. Furthermore, the epitope overlaps with the membrane proximal region (MPR) helices that sequester the fusion loop residues at the tip of domain I in the full-length prefusion conformation (**Fig. 4D**). This limited epitope accessibility on the native virion could explain the poor neutralizing activity of Fab 3-5 despite its high binding affinity for the prefusion-stabilized gB ectodomain construct. The prefusion specificity of Fab 3-5 is explained by the steric clashes between its heavy chain and domain I of the adjacent protomer in the postfusion conformation (**Fig. 4E**). Given that 3-5 is unable to bind postfusion gB trimers and is unlikely to bind prefusion membrane-anchored trimers, this antibody may have been elicited against an intermediate conformation of gB.

### Conservation and glycan shielding of HSV-2 prefusion-specific epitopes across human herpesvirus gBs

We analyzed the sequence similarity and glycan shielding of the prefusion-specific epitopes on HSV-2 gB targeted by mAbs 5-18 and 3-6 and compared these features across human herpesvirus gBs, focusing on the betaherpesvirus HCMV and the gammaherpesvirus EBV (**Fig. S15**). Sequence conservation analysis revealed that the quaternary epitope recognized by 5-18 is highly divergent among these viruses, with only 6 out of 44 contact residues conserved in HCMV and 7 out of 44 in EBV (**fig. S15A**). The epitope targeted by 3-6 also shows low conservation across these viruses, with 4 out of 24 residues conserved in HCMV and 8 out of 24 in EBV. Notably, the hinge region within this epitope is highly variable among human herpesvirus gBs, reflecting the divergent evolution of this site (**fig. S15B**). It is therefore unlikely that broadly reactive, pan-herpesvirus antibodies will be elicited against either site on gB.

In addition to sequence conservation, we also analyzed the glycan shielding of these epitopes. HSV-2 gB contains six N-linked glycosylation sites per protomer, four of which are resolved in our structures. Domains I and IV, which are major targets of neutralizing antibodies, lack glycosylation, and the quaternary epitopes recognized by 5-18 and 1-14 remain unshielded in HSV-2 gB (**fig. S15C**). An N-linked glycan at Asn671 is positioned near Fab 3-6, but this does not prevent the antibody from binding (**fig. S15C**). We next examined glycan shielding of prefusion epitopes on HCMV (*43*) and EBV (*44*) gB, for which glycans were modeled using Glycoshape (*45*) (**fig. S15C**). Structural analysis of the 5-18 contact sites on HCMV and EBV gB shows that, although the glycans at Asn586 and Asn555 in HCMV and Asn563 in EBV are positioned at the edge of the contact interface in domain IV, the major contact surfaces of the 5-18 epitope remain largely unshielded in both viruses (**fig. S15C**). In contrast, the 3-6 binding site, particularly the hinge region, contains a glycan in both HCMV (Asn341) and EBV (Asn290), suggesting glycan-mediated masking of a site that is highly vulnerable on HSV-2 gB (**fig. S15C**). Notably, this hinge-region glycan is conserved in HCMV, EBV, and HHV-6, but is absent from HSV-1, VZV, KSHV, and HHV-7, indicating divergent glycan-shielding strategies among human herpesviruses (**fig. S15B**).

### Prefusion-specific gB antibodies protect from HSV-2 challenge in a neonatal mouse model

Given the strong neutralizing activity of 3-6, 5-18, and 1-14, we assessed their ability to protect mice from a lethal challenge of HSV2-CNS11, a highly virulent clinical isolate. Briefly, 2-day-old C57BL/6J pups were administered 80 µg of mAb intraperitoneally (i.p), followed by an immediate intranasal (i.n.) challenge with 500 plaque-forming units (PFU) of HSV2-CNS11 (**Fig. 5**). Each of the three tested prefusion gB-specific mAbs provided statistically significant (p ≤ 0.002) partial protection at this treatment dose, as compared to the VRC01 control. All animals treated with the negative control HIV-1 envelope-directed mAb VRC01 failed to survive infection. At day 21 post-infection, the protection afforded by 3-6 was equivalent to that provided by HSV8 (gD-specific), with an overall survival rate of 91%. Treatment with 5-18 afforded 75% protection, which was comparable to but lower than the 87% protection provided by the gB-specific control mAb D48. Antibody 1-14 provided the least protection (20%). Collectively, these data demonstrate that mAb 3-6 can confer a high degree of protection against a lethal HSV2-CNS11 challenge in a neonatal mouse model, equivalent to that afforded by the gD-directed mAb HSV8.

**Fig 5.**
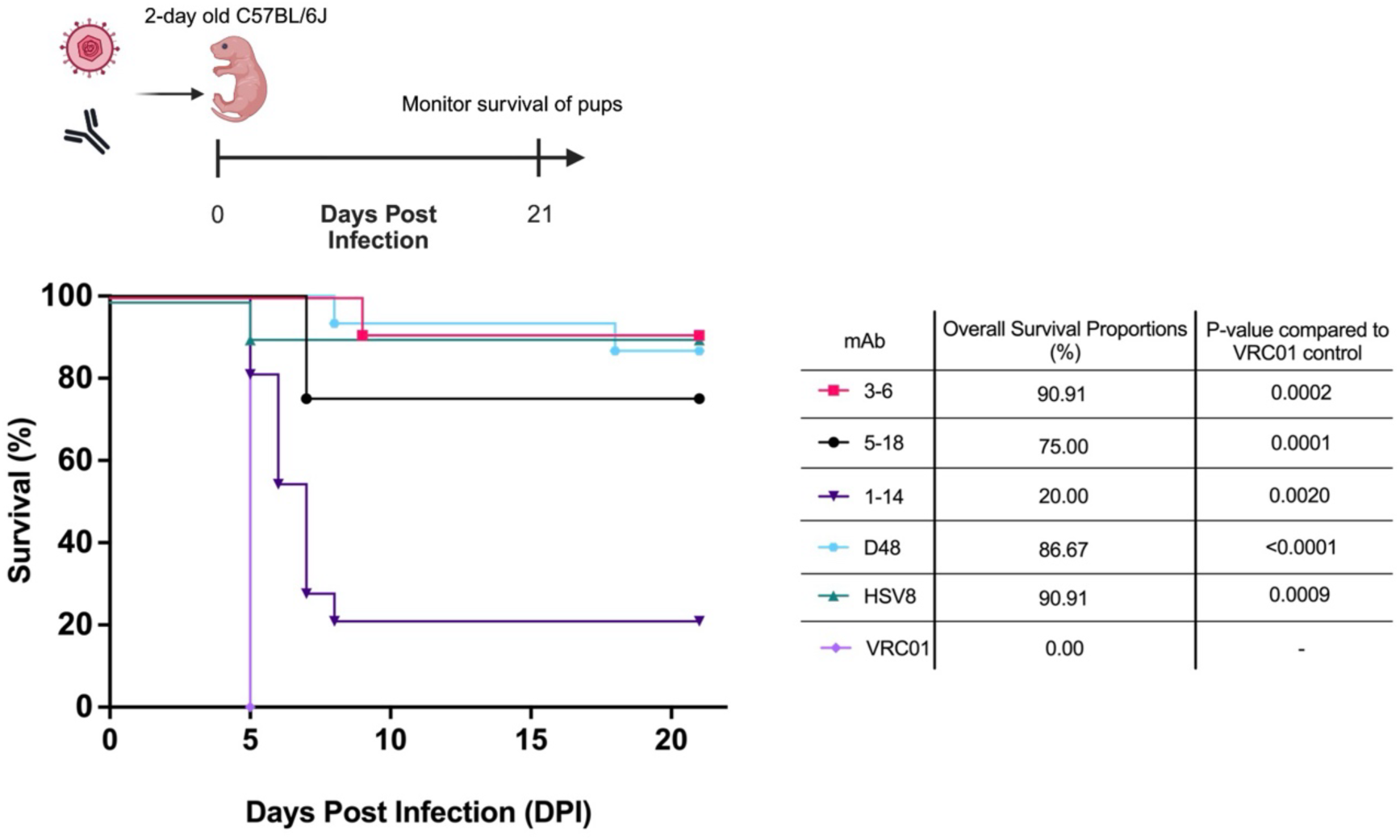
Prefusion-specific gB antibodies protect from viral challenge in a neonatal mouse model. Overall survival of 2-day old C57BL/6J mice infected intranasally with 500 plaque-forming units (PFU) of HSV-2-CNS11 and treated intraperitoneally with 80 μg monoclonal antibody IgG1 (3-6: n=11, 5-18: n=12, 1-14: n=15, D48: n=15, HSV8: n=11, VRC01: n=9) as shown on the experimental timeline (schematic created using Biorender.com). Statistical significance between the negative control VRC01 mAb and experimental antibodies was calculated by log(rank) (Mantel-Cox) test. Summary table with the overall survival proportions for each experimental group at 21 days post-infection as determined by the survival analysis Kaplan-Meier test.

## DISCUSSION

HSV is a widespread and clinically significant human pathogen that disproportionately affects vulnerable populations. Notably, HSV is an etiologic agent of neurologic and neonatal disease, often resulting in lifelong complications and substantial direct and indirect healthcare costs (*46, 47*). Preclinical studies have demonstrated that protective mAbs are highly effective for controlling perinatal HSV infection (*27, 29, 36, 37*); however, the structural and molecular basis of antibody recognition of the viral fusion glycoprotein B (gB)—particularly in its prefusion conformation—remains poorly defined. Here, we describe the identification of human antibodies that recognize epitopes exclusive to prefusion gB and potently cross-neutralize both HSV-1 and HSV-2. Two such antibodies, 5-18 and 3-6, conferred strong protection against lethal HSV-2 infection in a neonatal mouse model, further underscoring their therapeutic potential.

Prior to this work, prefusion-specific human antibodies targeting gB had not been characterized. This likely reflects both the substantial similarity between the exposed surfaces on prefusion and postfusion gB and the lack of prefusion-stabilized gB probes for B-cell sorting. Recently, a prefusion-specific camelid nanobody and a prefusion-specific murine antibody were isolated following immunization with vesicle-displayed native HSV-1 gB (*26*) or prefusion-stabilized HSV-1 gB (*25*), respectively; however, large-scale screening of human B cells specifically targeting prefusion gB has not been reported. Here, we address this limitation by applying LIBRA-seq with a prefusion-stabilized HSV-2 gB antigen (*39*) to PBMCs from HSV-seropositive donors, enabling the isolation of human prefusion-specific antibodies and the identification of previously uncharacterized vulnerable epitopes on HSV gB. With the recent development of soluble, prefusion-stabilized herpesvirus gB antigens and high-resolution structural information (*39–41*), there is now considerable potential to discover and develop additional potent prefusion gB-specific antibodies. Although prefusion-specific antibodies targeting HCMV and EBV gB may differ from those recognizing HSV-1 and HSV-2 gB due to distinct glycan shielding patterns (**fig. S15C**), it will be important to determine whether similarly potent neutralizing antibodies and discrete prefusion-specific epitopes exist across herpesvirus gB orthologs.

Antibodies 5-18 and 3-6 potently neutralized both HSV-1 and HSV-2, with 3-6 demonstrating the greatest potency against both clinical isolates (**Fig. 2D-E**). When co-administered with a lethal dose of HSV2-CNS11 in a neonatal mouse study, mAb 3-6 conferred protection comparable to previously characterized neutralizing antibodies HSV8 (gD-specific) and D48 (gB-specific), highlighting its therapeutic potential (**Fig. 5**). Given the multistep nature of herpesvirus entry (*48*), therapeutic strategies employing antibody cocktails that target multiple epitopes within a single glycoprotein or span distinct viral glycoproteins may provide enhanced protection. Because gD mediates viral attachment (*49*) and gD-directed antibodies have been extensively studied and have shown robust neutralizing activity (*28*), combining gD-specific antibodies with prefusion gB-specific antibodies may represent an improved strategy for HSV prevention or treatment. Such combinations would target sequential steps in viral entry and may yield synergistic antiviral effects.

In these studies, we determined that the prefusion-specific antibody 3-5 binds a membrane-proximal epitope on gB. Structural analysis indicates that 3-5 is unlikely to engage prefusion gB in the viral membrane, as its predicted angle of approach would result in steric clashes with the lipid bilayer and MPR helices. Consistent with this interpretation, 3-5 exhibits poor neutralizing activity against HSV-2, suggesting that it may recognize an intermediate conformational state of gB, such as after domain I has disengaged from the MPR but before insertion of the hydrophobic fusion loops into the target cell membrane. Intermediate conformations of class III fusion proteins have been visualized by cryo-electron tomography, including for vesicular stomatitis virus G (*50*). Analogously, non-neutralizing prefusion-specific antibodies have been described for HIV-1, where antibodies targeting membrane-proximal gp41 base epitopes are preferentially elicited, thereby constraining the induction of broadly neutralizing responses (*51, 52*). Recent studies further indicate that prefusion-stabilized HSV-2 gB elicits immune responses comparable to those induced by postfusion gB (*39*), underscoring the challenge of selectively focusing humoral immunity on functionally vulnerable prefusion epitopes. Systematic characterization of B cell lineages, coupled with antigen-design strategies such as glycan shielding or epitope masking, can be utilized to preferentially enhance the activation of B cells encoding potently neutralizing and protective antibodies against HSV. Furthermore, the results from these studies suggest that potently neutralizing mAbs targeting the prefusion form of gB represent an attractive therapeutic option for preventing HSV-associated disease in vulnerable populations.

## ACKNOWLEDGMENTS

We thank all members of the Georgiev laboratory and McLellan laboratory for their support and feedback. We thank David Flaherty, Olivia Murfield, Emma McLaughlin, and Brittany Matlock from the VUMC Flow Cytometry Shared Resource for their help with cell sorting. The VUMC Flow Cytometry Shared Resource is supported by the Vanderbilt Ingram Cancer Center (P30 CA68485) and the Vanderbilt Digestive Disease Research Center (DK058404). We thank Angela Jones, Jamie Roberson, and Latha Raju with the Vanderbilt Technologies for Advanced Genomics Core (VANTAGE) for providing technical assistance with library production and sequencing. VANTAGE is supported in part by CTSA (5UL1 RR024975-03), the Vanderbilt-Ingram Cancer Center (P30 CA68485), the Vanderbilt Vision Center (P30 EY08126), and NIH/NCRR (G20 RR030956). We thank Dr. Axel Brilot and Dr. Evan Schwartz at the Sauer Structural Biology Laboratory at UT Austin for assistance with cryo-EM data collection, which is supported in part by The University of Texas at Austin College of Natural Sciences and award RR160023 of the Cancer Prevention and Research Institute of Texas. The funders had no role in the conceptualization or execution of any studies or drafting of the manuscript.

National Institutes of Health grant R01AI175245 (ISG, PA, GJ, PTW, AAA)

National Institutes of Health grant R01AI176646 (NNM, DAL, MEA)

National Institutes of Health grant R01DK131070 (RHB)

Welch Foundation grant number F-0003-19620604 (JSM)

National Institutes of Health fellowship NIH F31DK141224 (LEB)

## AUTHOR CONTRIBUTIONS

Conceptualization: PA, JK, NVJ, AAA, JSM, ISG

Investigation: PA, JK, NNM, PTW, GJ, LEB, NVJ

Supervision: MEA, DAL, RHB, JSM, ISG

Writing – original draft: PA, JK, JSM, ISG

Writing – review & editing: PA, JK, NNM, PTW, LEB, GJ, AAA, MDS, NVJ, RHB, DAL, MEA, JSM, ISG

## DATA, CODE, AND MATERIALS AVAILABILITY

All data are available in the main text or the supplementary materials.

## MATERIALS AND METHODS

### Cells and viruses

FreeStyle 293F cells (Gibco), Expi293 cells (Gibco), Vero cells (ATCC, cat. no. CCL-81), HEp-2 cells (ATCC, cat. no. CCL-23), and HEK293T cells (ATCC, cat. no. CRL-3216) were cultured according to vendor instructions. Virus strains used in this study included HSV2-G (*53*), HSV1-NS (*54*), and HSV2-CNS11 (*55*). For virus propagation, Vero cells were used to prepare viral stocks as described previously (*56, 57*).

### Mice

C57BL/6J (RRID:IMSR_JAX:000664) mice were either purchased from The Jackson Laboratories or bred in animal facilities at Dartmouth College in accordance with Institutional Animal Care and Use Committee protocols (Dartmouth College IACUC protocol 2151).

### Human blood sample collection and preparation

Prospective blood collection and peripheral blood mononuclear cell (PBMC) isolation from HSV-2 positive donors were procured from BioCollections Worldwide, Inc. (Miami, FL) and certified by BioPlex 2200 HSV-1 & HSV-2 IgG kit on the BioPlex 2200 System. Healthy donor PBMCs with no recorded history of prior infection were purchased commercially from StemCell Technologies (catalog number 70025.3). Samples arrived frozen and were stored at -135 °C until use.

### HSV gB expression and purification

HSV-1 gB ectodomain (Strain KOS; Uniprot: P06437, residues 31–725) was engineered in both postfusion and prefusion-stabilized forms using the same prefusion-stabilizing substitutions as the HSV-2 gB constructs (Strain HG52; Uniprot: P06763, residues 23–731; postfusion gB-B1 and prefusion-stabilized gB-O3)(*58*). For HSV-1 prefusion gB, the N-terminal disordered region (residues 32–78) was truncated to improve expression, an interprotomer disulfide bond (T243C/E679C) and two proline substitutions (S688P and T669P) were introduced to stabilize the prefusion conformation, and substitutions H172N and Y174T were incorporated to reduce aggregation (**fig. S5**). All constructs were transiently expressed in FreeStyle 293F cells using polyethylenimine (PEI) transfection reagent. Cultures were maintained at 37 °C and 8% CO₂, and the media were harvested 6 days post-transfection. The supernatants were filtered through a 0.22 µm membrane and loaded onto Strep-Tactin 4Flow resin (IBA Lifesciences) by gravity. After washing with 100 mM Tris-HCl pH 8.0, 150 mM NaCl, and 1 mM EDTA, the bound proteins were eluted with 100 mM Tris-HCl pH 8.0, 150 mM NaCl, 1 mM EDTA, and 50 mM biotin (IBA Lifescience). The elution fractions were incubated overnight at 4 °C with 5% (w/w) His-tagged HRV3C protease to remove purification tags. The samples were then purified by size-exclusion chromatography (SEC) using a Superose 6 Increase 10/300 GL column (Cytiva) equilibrated in phosphate-buffered saline (PBS).

### Oligonucleotide barcodes

Oligonucleotides composed of a 15 bp antigen barcode were used with sequence modifications to anneal to the template switch oligo on the 10x bead-delivered oligos and truncated TruSeq small RNA read 1 sequence in the following structure: 5′ CCTTGGCACCCGAGAATTCCANNNNNNNNNNNNNCCCATATAAGA∗A∗A-3′, where N represent the antigen barcode. For each antigen, we used the following barcode sequences: HSV-2 gB-G3 prefusion (GACAAGTGATCTGCA), HSV-2 gB B1 postfusion (AACCCACCGTTGTTA), HCMV gB C7 prefusion (CAGTAGATGGAGCAT), HCMV gB base postfusion (ATCGTCGAGAGCTAG), EBV gB C3-GT prefusion (TGTGTATTCCCTTGT), EBV gB base postfusion (GCTCCTTTACACGTA), HIV BG505 gp140 V9.3 SOSIP (TCACAGTTCCTTGGA), and influenza HA NC99 (GGTAGCCCTAGAGTA). Oligos were ordered from Sigma-Aldrich and IDT with a 5′ amino modification and HPLC purified.

### Antigen labeling with DNA oligonucleotide barcodes

For each antigen, the unique DNA barcodes were directly conjugated using SoluLINK Protein-Oligonucleotide Conjugation kit (Vector Labs, S-9011) according to the kit’s protocol. Each antigen was biotinylated using EZ link (Thermo), desalted and modified with S-HyNic. The 4FB-modified DNA barcodes were directly conjugated to the biotinylated antigen with S-HyNic to create a stable bond between the protein and oligonucleotide. The antigen-oligonucleotide conjugate concentrations were determined using a bicinchoninic acid (BCA) assay, and the HyNic molar substitution ratios of each antigen-oligonucleotide conjugate were determined using a NanoDrop according to the SoluLINK protocol instructions. Excess oligonucleotides were removed from the protein-oligonucleotide conjugates using an AKTA FPLC and were subsequently verified using SDS-PAGE and silver stain. The optimal amounts of antigen-oligonucleotide conjugates to be used in antigen-specific B cell sorting were then determined through flow cytometry titration experiments on cell lines expressing BCRs of known specificities.

### Antigen-specific B cell sorting

PBMCs were thawed and counted using Trypan Blue viability stain. Cells were washed with a solution of Dulbecco’s phosphate-buffered saline (DPBS) supplemented with 0.1% bovine serum albumin (BSA). Samples were resuspended in DPBS-BSA and stained with cell markers: CD14 (BD, cat no. 301819, clone M5E2, 1:10), Ghost Dye Red 780 Viability (Cell Signaling Technology, cat no. 18452S, 1:1000), CD3 (Cytek Biosciences, cat no. 35-0037, clone OKT3), CD19 (BD, cat no. 563036, clone SJ25C1), and IgG (BD, cat no. 551497, clone G18 145). Samples were resuspended in DPBS-BSA before surface staining for 30 min. Additionally, antigen-oligonucleotide conjugates were incorporated into the stain. After 30 min incubation in the dark at room temperature, cells were washed three times with DPBS-BSA at 300g for 5 min. Next, the samples were incubated for 15 min at room temperature with Streptavidin-PE (Invitrogen, cat no. S866) to label cells with bound antigen. The cells were again washed three times with DPBS-BSA, resuspended in DPBS, and sorted by FACS. Antigen-positive B cells were bulk sorted and sequenced at Vanderbilt Technologies for Advanced Genomics (VANTAGE) at an appropriate target concentration for 10X Genomics library preparation and subsequent sequencing. FACS data were analyzed using FlowJo (BD Biosciences).

### Sample and library preparation for sequencing

Single-cell suspensions were processed using the Chromium Controller microfluidics device (10X Genomics) and B cell Single Cells V(D)J solution as per the manufacturer’s instructions. The aim was to capture 10,000 B cells per 1/8 10X cassette. Slight modifications were made to intercept, amplify, and purify the antigen barcode libraries, as previously described (*59*).

### Sequence processing and bioinformatics analysis

For bioinformatic processing of the LIBRA-seq sequencing reads, we followed our established pipeline, which takes paired-end FASTQ files of oligonucleotide libraries as input. This pipeline processes and annotates reads for cell barcodes, unique molecular identifiers (UMIs) and antigen barcodes, resulting in a cell-barcode and antigen-barcode UMI count matrix (*38*). B cell receptor contigs and antigen-barcode libraries were processed using CellRanger 3.1.0 (10x Genomics) with GRCh38 Human V(D)J 7.0.0 as reference. From the BCR sequencing, 8,467 functional paired heavy- and light-chain sequences were recovered. B cell receptor contigs were aligned to IMGT reference genes using High-V Quest (*60*).

### Antibody expression and purification

Variable heavy and light genes were inserted into custom plasmids that encode the constant region for the human IgG1 heavy chain and respective lambda and kappa light chains (Genscript). The antibodies were expressed in Expi293 cells by co-transfecting heavy chain and light chain expression plasmids using PEI or Expifectamine transfection reagent (Thermo Fisher Scientific) and cultured for 4–5 days. Cultures were harvested and centrifuged, and supernatant was filtered through 0.45 μm Nalgene Rapid Flow Disposable Filter Units with PES membrane. The filtered supernatant was run over a column containing Protein A agarose resin that was equilibrated with PBS. The column was washed with PBS, and then the antibodies were eluted with 100 mM glycine-HCl at pH 2.7 directly into a 1:10 volume of 1 M Tris-HCl pH 8.0. Eluted antibodies were buffer exchanged into PBS using Amicon Ultra centrifugal filter units, centrifuging and topping off three times with PBS, and finally concentrated. The antibodies were analyzed by SDS-PAGE. Antibody plasmids were sequenced to confirm the expected heavy and light chains.

### Indirect enzyme-linked immunosorbent assay (ELISA)

For indirect ELISA assays, maxisorp 96-well flat-bottomed plates (Thermo Fisher Scientific) were coated with recombinant gB antigen at 2 μg mL^−1^ in PBS overnight at 4 °C. Subsequent washes were performed with PBS-T (PBS supplemented with 0.05% v:v Tween20 (Millipore-Sigma)). Blocking was performed in 5% BSA in PBS-T for 2 h at room temperature and then washed three times with PBS-T. Primary antibodies were serially diluted 1:5 in 1% BSA in PBS-T with a starting concentration of 10 μg mL^−1^ and incubated at room temperature for 1 h, washed, and detected via a horseradish peroxidase-conjugated anti-mouse IgG secondary diluted 1:10,000 (Jackson ImmunoResearch). One-step TMB-ELISA substrate solution (Thermo Fisher Scientific) was added to develop for 5 min and quenched with 1 N sulfuric acid. Absorbance at 450 nm (A_450_) was measured on a microplate reader. ELISAs were performed in technical and biological duplicates.

### Antibody competition ELISA

For competition ELISA assays, maxisorp 384-well microtiter plates (Thermo Fisher Scientific) were coated with recombinant gB antigen at 2 μg/mL in PBS overnight at 4 °C. Subsequent washes were performed three times with PBS-T. Blocking was performed in 5% BSA in PBS-T for 2 h at room temperature, followed by three washes with PBS-T. Primary competitor antibodies at 10 μg/mL were added to wells incubated for 1 h at room temperature. Without washing off the competitor antibody, a biotinylated preparation of antibodies was added to wells with primary antibody at a concentration of 1 μg/mL and incubated for 1 h at room temperature. Plates were washed three times with PBS-T and bound antibodies were detected via a horseradish peroxidase-conjugated mouse anti-biotin secondary diluted 1:4,000 (SouthernBiotech). One-step TMB-ELISA substrate solution (Thermo Fisher Scientific) was added, incubated for 5 min, and quenched with 1 N sulfuric acid. Absorbance at 450 nm (A_450_) was measured on a microplate reader. The A_450_ signal obtained for binding of the biotin-labelled reference antibody in the presence of the unlabeled tested antibody was expressed as a percentage of the binding of the reference antibody alone. Competition ELISAs were performed in technical and biological duplicates.

### Prefusion specificity measurement

Bio-layer interferometry (BLI) experiments were conducted on an OctetRED96e instrument (Sartorius) at 21°C with a shaking speed of 1000 rpm. For determination of prefusion specificity, HSV-2 gB-specific IgG antibodies were immobilized onto anti-human IgG Fc Capture (AHC) biosensors (Sartorius) to a response level of approximately 1.0 nm in HBS-EP buffer (10 mM HEPES pH 7.4, 150 mM NaCl, 3 mM EDTA, 0.05% v/v surfactant P20) supplemented with 0.1 % w/v BSA. Loaded sensors were then dipped into wells containing 200 nM of HSV-2 or HSV-1 prefusion-stabilized gB or postfusion gB for 180 s (association phase), followed by 180 s of dissociation. Reference sensors were incubated in buffer alone under identical conditions. Data were reference-subtracted and analyzed using the Octet Analysis Studio (v13.0, Sartorius).

### Public antibody analysis

Prefusion-specific gB mAbs (1–14, 5–18, 3–5, 3–6) and prefusion-preferring gB mAbs (1–13, 5–13) were compared against a database of previously published antibodies consisting of 1.7 million paired antibody sequences. This database consists of a previously curated dataset (*61*), expanded with antibodies from public databases (*62–69*). In comparing antibody pairs, V gene paralogues were treated as the same, and CDR3 identity was calculated by employing the Levenshtein distance metric, measuring the difference between amino acid sequences and dividing it by the length of the longer CDR3 region in the pair to obtain sequence identity. Scatterplots in **fig. S3A** depict these pairs of antibodies.

### Affinity measurement

For kinetics analyses, HSV-2 IgGs containing an HRV3C protease cleavage site engineered between the CH1 and CH2 domains of the heavy chain were incubated with 5% (w/w) His-tagged HRV3C protease at 4°C overnight. The reaction mixtures were then passed over Protein A and Ni-NTA resins to remove the cleaved Fc fragment and excess protease. The resulting Fabs were purified by SEC on a Superdex 200 Increase column (Cytiva) equilibrated in PBS. Purified soluble HSV-2 and HSV-1 prefusion-stabilized gB bearing a C-terminal 8xHis-tag were immobilized onto Ni-NTA biosensors (Sartorius) to a response level of approximately 0.7–0.8 nm in HBS-P buffer (10 mM HEPES pH 7.4, 150 mM NaCl, 0.005% v/v Surfactant P20) with 20 mM imidazole and 0.1% w/v BSA. After establishing a 180s baseline, the immobilized HSV-2 and -1 gB sensors were dipped into wells containing two-fold serial dilutions of Fabs. Sensors immobilized with gB proteins were run in parallel using buffer-only as a reference control. Dissociation was monitored for 600 s by transferring sensors into buffer-only wells. All data were reference-subtracted and globally fit to a 1:1 binding model using the Octet Analysis Studio (v13.0, Sartorius).

### Cryo-EM sample preparation and data collection

Purified HSV-2 prefusion-stabilized gB was mixed in PBS to a final concentration of 4.0 mg mL^−1^ with 1.2-fold molar excess of Fabs 5-18, 1-14, or a combination of 3-5 and 3-6. Complexes were incubated for 30 min at 4 °C before adding 10X CMC CHAPS (VitroEase™ Buffer Screening Kit, Thermo Fisher Scientific) to a final concentration of 0.25X CMC. Immediately after CHAPS addition, 3.0 μL of sample was applied to UltrAuFoil R1.2/1.3 grids (Electron Microscopy Sciences) that had been glow-discharged using a PELCO easiGlow (Ted Pella) for 30 s at 20 mA. Grid vitrification was performed using a Vitrobot Mark IV (Thermo Fisher Scientific) operated at 4 °C in 100% humidity. A blot force of 8 was applied for 6 s to remove excess liquid before plunge-freezing the grids into liquid ethane. For data collection, 2,553 movies for the Fab 1-14 complex, 2,551 movies for the 5-18 Fab complex, and 3,060 movies for the Fab 3-5/Fab 3-6 complex were collected from single grids using a Glacios TEM (Thermo Fisher Scientific) equipped with a Falcon 4 detector (Thermo Fisher Scientific). Data for the 1-14 and 1-15 complexes were acquired at a 30° stage tilt, whereas data for the 3-5/3-6 complex were collected without tilt. All movies were recorded using SerialEM v4.0.10 automation software (*70*). Movies were acquired at a calibrated pixel size of 0.933 Å/pixel (total dose: 49 e⁻/Å²). Additional details about data collection parameters are summarized in **Table S4.**

### Cryo-EM processing and structure determination

Motion correction, CTF estimation, and particle picking were performed using CryoSPARC Live. All subsequent image processing steps were performed in CryoSPARC v4.7.1 and v5.0.0-privatebeta.3 (*71*). Briefly, particles underwent two rounds of *ab initio* reconstruction and heterogeneous refinement, followed by re-extraction at 384 px. Multiple rounds of non-uniform refinements with optimized parameters were then performed to improve resolution, followed by reference-based motion correction (**fig. S8, fig. S9 and fig. S12**). For the Fabs 3-5 and 3-6 complex, 3D classification was performed (**fig. S12**). Final map sharpening was performed using EMReady v2.2.2 (*72*) within the SBgrid environment (*73*). Initial models were generated by combining the HSV-2 prefusion-stabilized gB structure (*39*) with Alphafold3-predicted models of the variable regions of 5-18, 1-14, 3-5, and 3-6, produced by inputting their VH and VL sequences (https://alphafoldserver.com) (*74*). The highest-confidence output model was docked into the refined cryo-EM map via ChimeraX (*75*). Missing residues were manually built in Coot (*76*), and structures were interactively refined using a combination of Phenix (*77*), Coot, and ISOLDE (*78*). N-acetyl-glucosamine (NAG) and β-mannose (BMA) were added to glycosylation sites and refined using ChimeraX and ISOLDE. Antigen-antibody interfaces and buried surface areas were analyzed using the PDBe PISA service (European Bioinformatics Institute; http://www.ebi.ac.uk/pdbe/prot_int/pistart.html) and ChimeraX. Secondary structure elements were derived from the HSV-2 gB structure in complex with Fabs 3-6 and 3-5, and plotted alongside sequence alignments of HSV-1 and HSV-2 gB generated using the UniProt Clustal Omega server (*79*).

### Plaque reduction neutralization test

Serial dilutions of mAbs were incubated with 75–160 plaque-forming units (PFU) of HSV1-NS or 65–120 PFU of HSV2-G for 1 h at 37 °C and 5% CO_2_. This mixture was added to a confluent layer of Vero cells in 6-well plates and incubated for 1 h at 37 °C and 5% CO_2_, with shaking every 15 minutes, after which a 1:1 methylcellulose:media overlay was added. The plates were then incubated for 44 to 48 h at 37 °C and 5% CO_2_. Cells were fixed with a 1:1 ethanol:methanol solution for 30 min prior to staining with 12% Giemsa for 24 to 48 h. The stain was then removed, and plaques were counted to determine the extent of neutralization relative to virus-only controls.

### Intracellular and extracellular flow cytometric analysis

HEp-2 and HEK293T cell lines were harvested for flow cytometry staining. Cells were incubated with Alexa Fluor 700 NHS ester (succinimidyl ester) (Thermo Fisher Scientific) prior to fixation/permeabilization to enable viability assessment. For extracellular staining, cells were incubated on ice with 25 µg/mL unlabeled experimental and control IgG mAbs diluted in flow cytometry staining buffer, or flow cytometry staining buffer alone (secondary controls). Cells were washed, and rat anti-human IgG Fc-PE/Cy7 (Biolegend cat no. 410721, clone M1310G05, 1:1500) was used to detect human IgG mAbs. For intracellular staining, cells were fixed and permeabilized using the BD Cytofix/Cytoperm kit, followed by incubation with 25 µg/mL experimental or control mAbs diluted in the kit perm/wash buffer per kit instructions. Cells were washed, and rat anti-human IgG Fc-PE/Cy7 (Biolegend cat no. 410721, clone M1310G05, 1:1500) was diluted in perm/wash buffer and used to detect human IgG mAbs. All cells were resuspended in 1% paraformaldehyde, 2 mM EDTA in PBS and stored at 4 °C prior to acquisition on a BD Biosciences LSR Fortessa flow cytometer. Data were analyzed using FlowJo software (BD Biosciences). Gating strategies and IgG^+^ plots can be found in **fig. S7.**

### Survival study in a neonatal mouse model

Two-day-old C57BL/6J pups under 1% isoflurane anesthesia were treated with 80 µg of mAb in a 20 µL volume intraperitoneally (i.p.) using a 25 µL Hamilton syringe and then immediately challenged intranasally (i.n.) with 500 PFU of HSV2 CNS11 in a 5 µL volume and monitored for survival for 21 days post-infection. Survival endpoints were defined in accordance with the criteria specified in the IACUC protocol (excessive morbidity such as hunching, spasms and/or paralysis and/or >10% weight loss from the previous measurement). All animal procedures were performed following approval by Dartmouth’s Institutional Animal Care and Use Committee and in accordance with Dartmouth’s Center for Comparative Medicine and Research policies.

### Statistical methods

All statistical analyses were performed using GraphPad Prism version 9.0.0. Results of the challenge study were graphed as a cumulative Kaplan-Meier survival curve, and comparisons between groups were performed using a log-rank (Mantel-Cox) test. *In vitro* neutralization curves were generated by fitting data points using a variable slope and a four-parameter regression curve (best-fit method). ELISA data were analyzed using variable slope and a four-parameter regression curve, with area under the curve (AUC) reported from log-transformed values. The number of replicates per experiment is indicated in the figure legends. No data or experiments were excluded unless there were technical issues with the experiment, and outliers were not excluded. Exact P values are denoted in the figures.

## SUPPLEMENTAL FIGURES

**Fig. S1.**
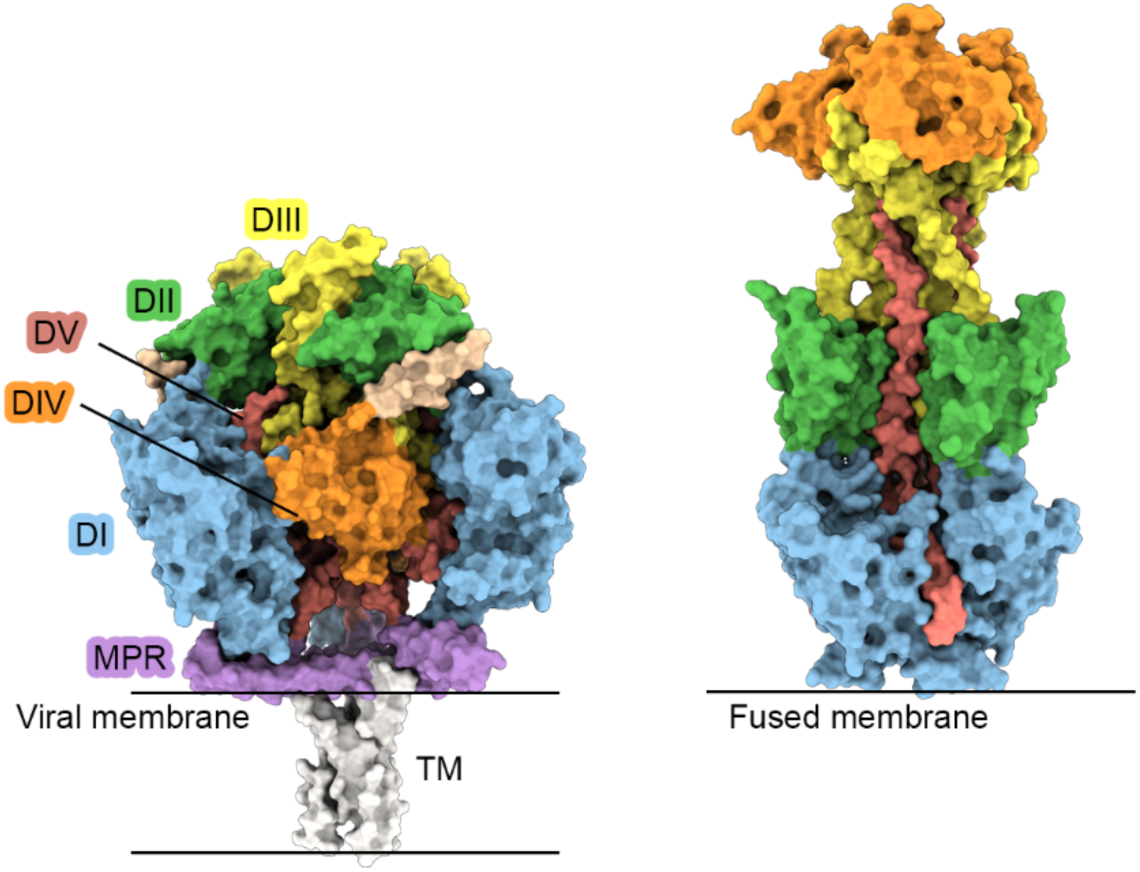
Structural comparison of prefusion and postfusion gB. Side views of prefusion (PDB ID: 9Q9L) and postfusion (PDB ID: 9IH8) HSV-2 gB are shown as molecular surfaces. Domain (D) I is colored blue, DII is colored green, DIII is colored yellow, DIV is colored orange, DIV is colored red, and

**Fig. S2.**
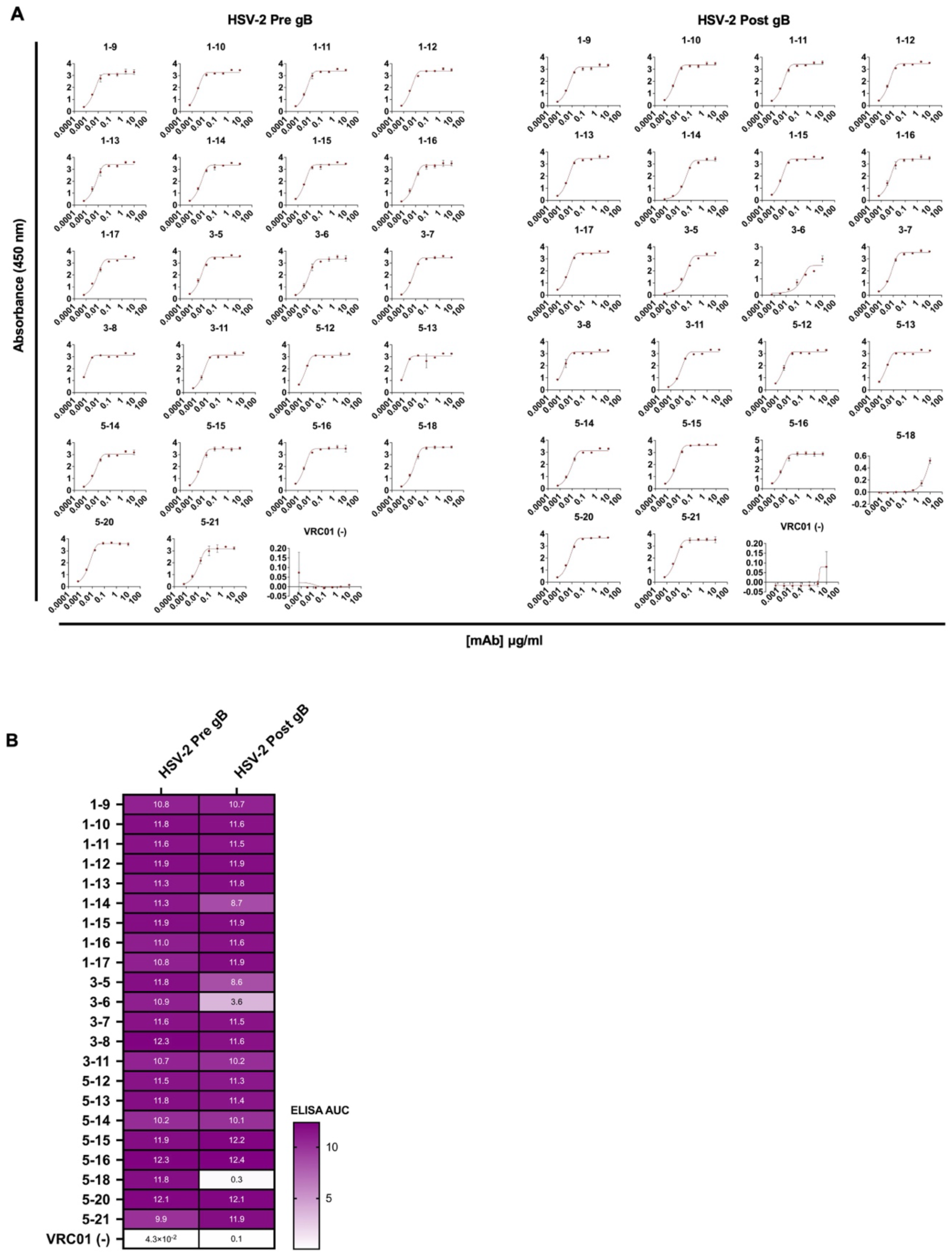
Antibodies discovered by LIBRA-seq bind to HSV-2 pre- and postfusion gB. **(A)** Indirect ELISA absorbance measurement of recombinantly produced antibodies binding to HSV-2 prefusion or postfusion gB. **(B)** Area under the curve (AUC) of transformed absorbance values. Experiments were performed in technical and biological duplicates.

**Fig. S3.**
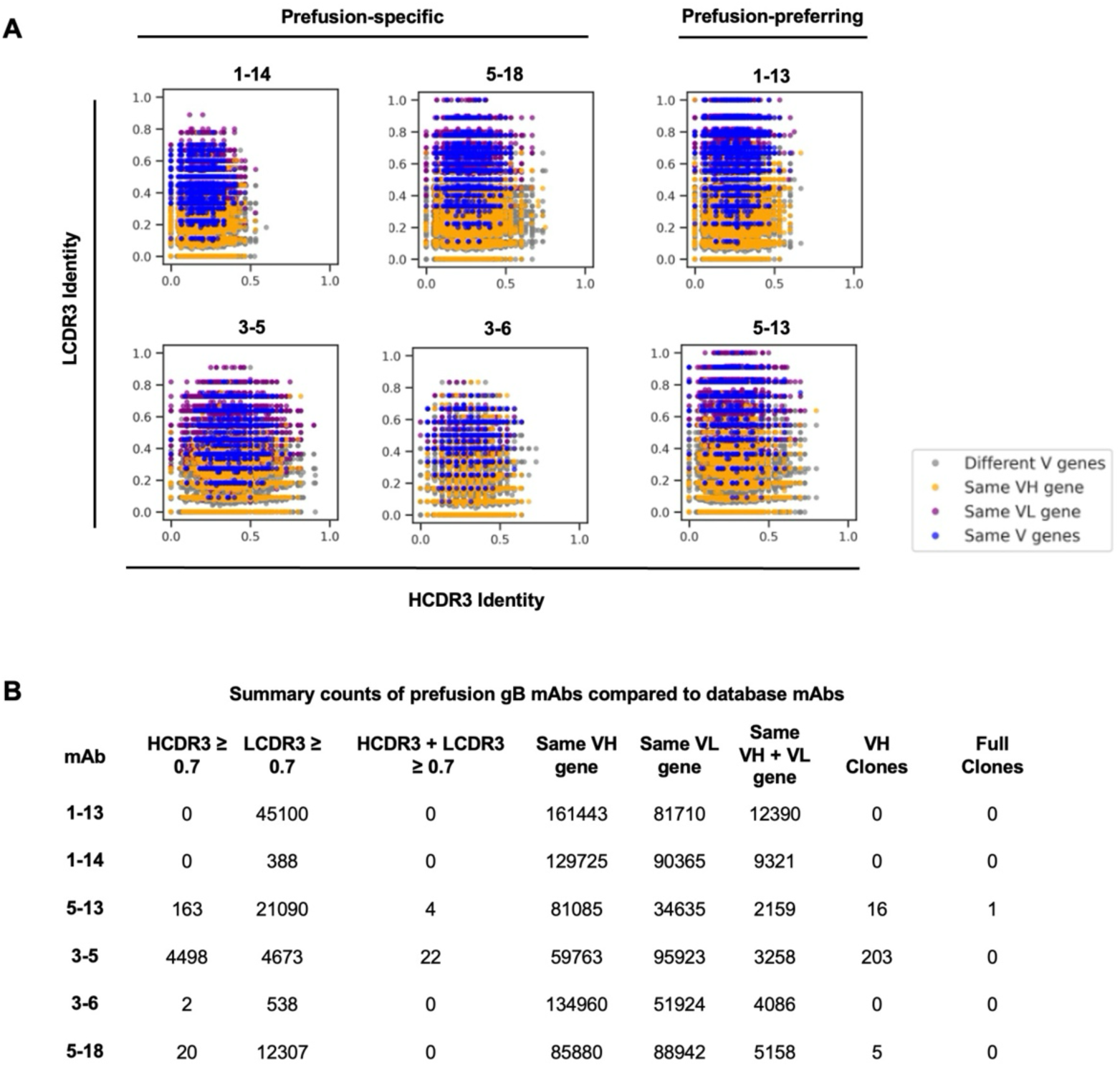
Comparison of prefusion gB mAbs to published antibodies. **(A)** Sequence similarity of prefusion gB mAbs to a public database of 1.7 million paired antibody sequences. Each point represents an antibody from the database, compared with the prefusion gB mAb (represented by the title for each panel). Characterized by their identities in the CDR3 of the heavy chain (represented on the x-axis) and the light chain on the y-axis. The color coding of each dot in the plots corresponds to the matching of V genes between pairs of antibodies: Blue dots indicate identical use of both VH and VL genes; orange signifies matching VH gene use; purple represents matching VL gene use; and gray denotes cases in which neither VH nor VL genes match. **(B)** Shown are the counts based on comparisons shown in panel **A**. Full clones are defined as having matching V-gene usage and ≥ 0.7 CDR3 identity.

**Fig. S4.**
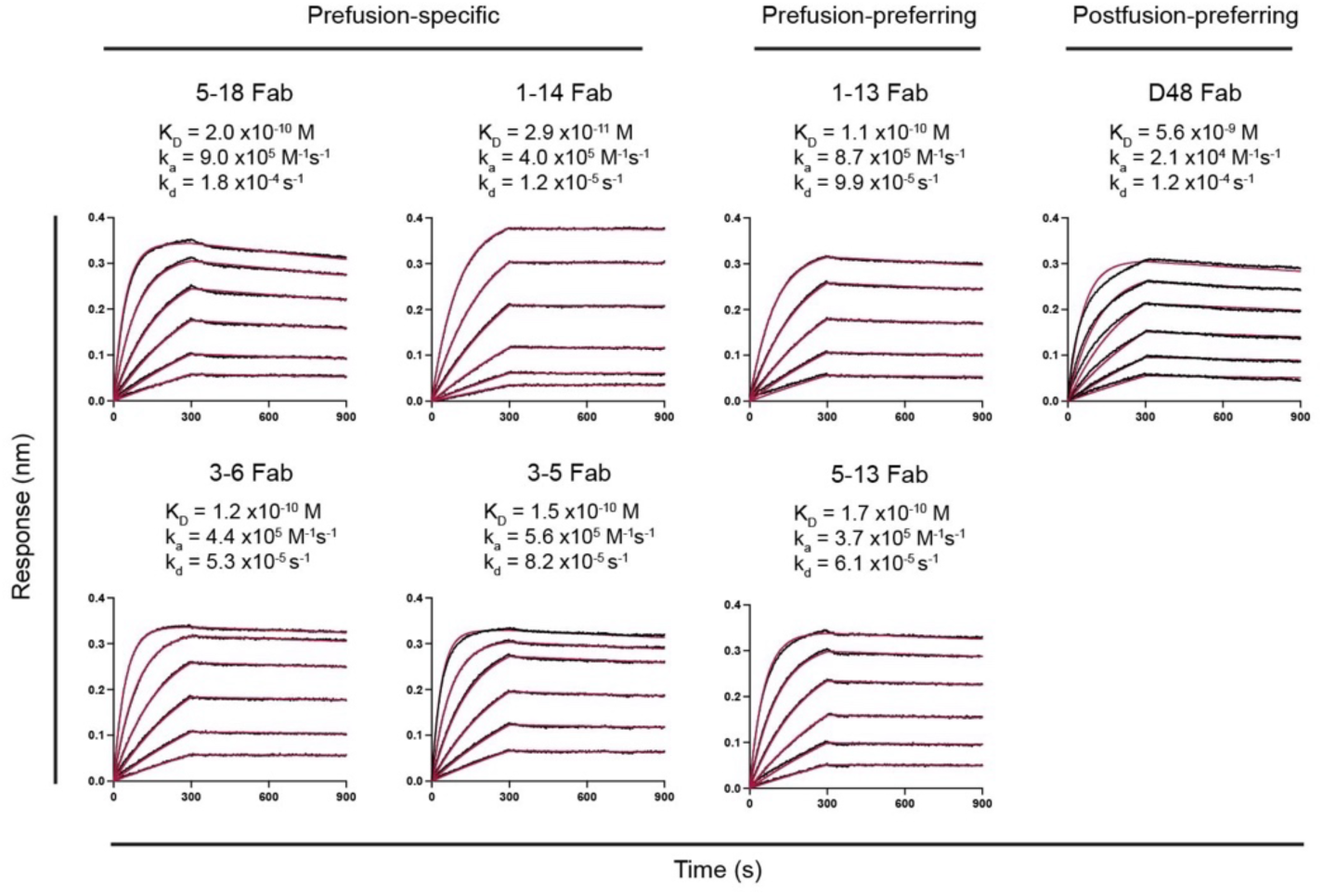
Kinetics of isolated Fabs binding to HSV-2 prefusion stabilized gB. BLI sensorgrams for the association and dissociation kinetics of prefusion-specific and prefusion-preferring Fabs, as well as the postfusion-preferring control Fab D48, binding to HSV-2 prefusion-stabilized gB. Experimental data (black) were globally fit to a 1:1 binding model, with the fitted curves shown in red. The calculated values for *k_d_*, *k_a_*, and *k_d_* are listed above each curve.

**Fig. S5.**
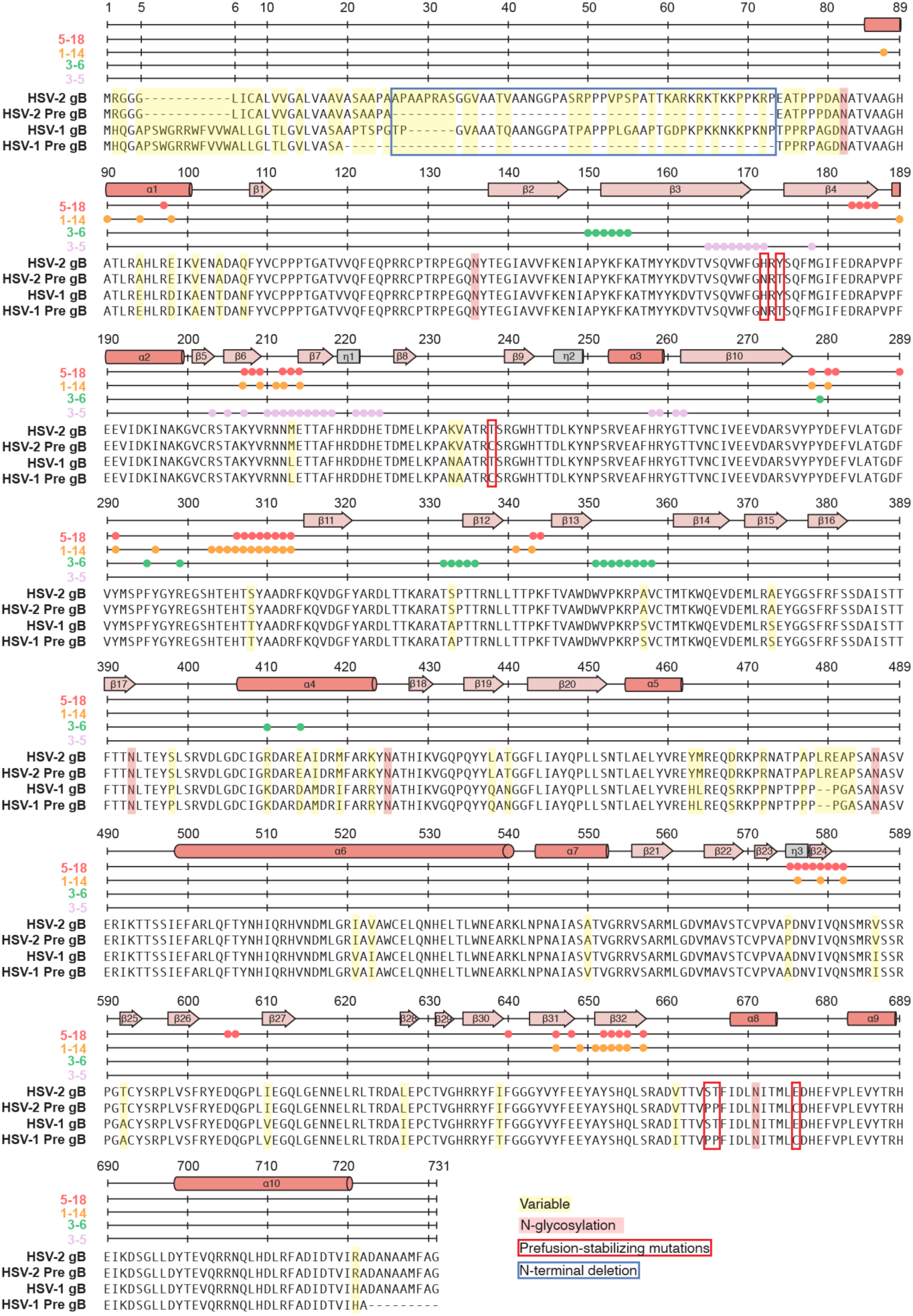
Sequence alignment of HSV-2 and HSV-1 gB. Sequence alignment of HSV-2 gB ectodomain residues 1–731 and HSV-1 gB ectodomain residues 1–725. Residues involved in antibody interactions are indicated by colored circles above the alignment. Secondary structure elements were derived from the coordinates of HSV-2 gB bound to Fabs 3-5 and 3-6 and are annotated above the alignment. Variable positions between HSV-2 and HSV-1 gB are highlighted in yellow, *N*-linked glycosylation sites are in red, prefusion-stabilizing substitutions are annotated by red boxes, and the engineered N-terminal deletion is indicated by a blue box.

**Fig. S6.**
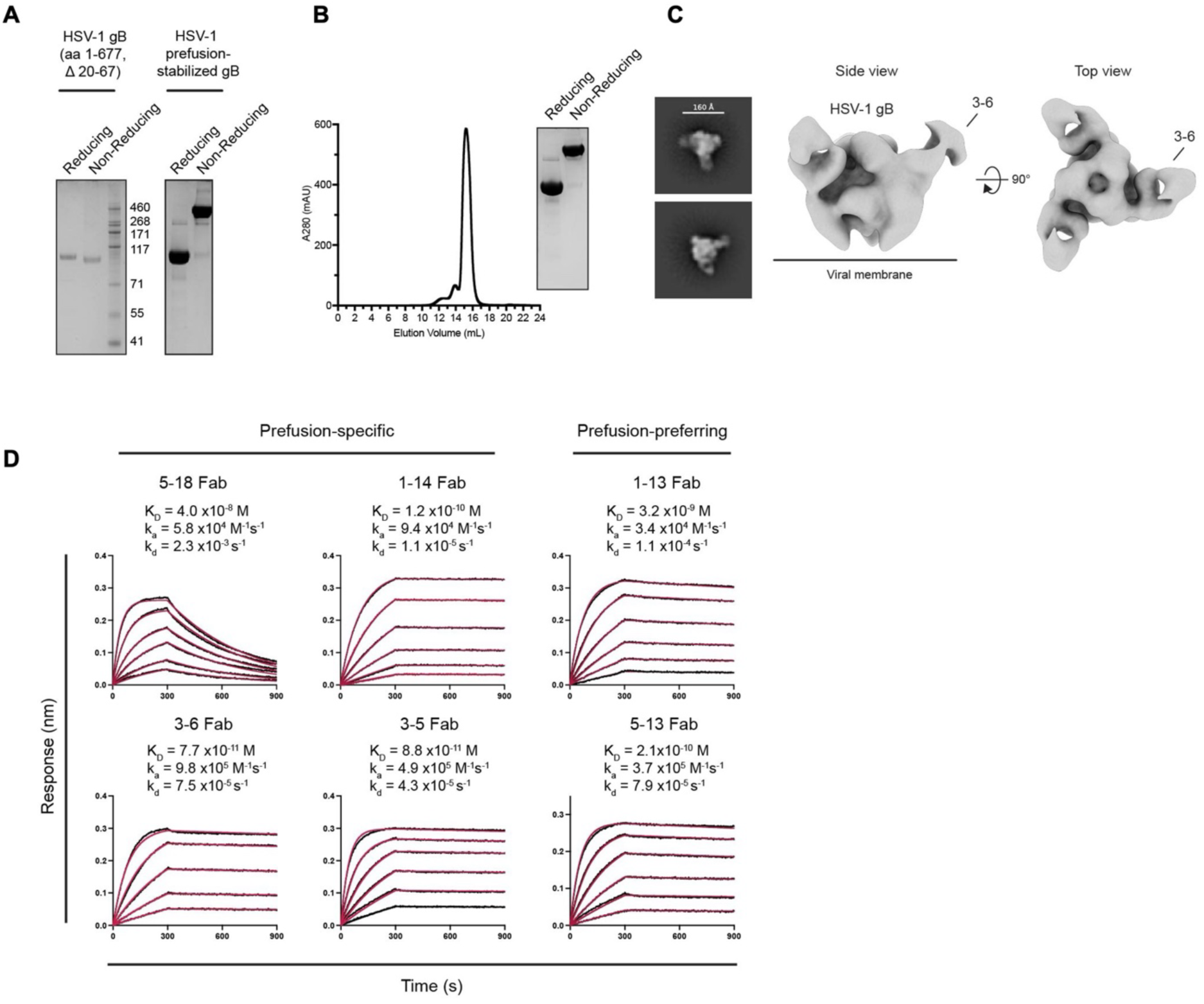
Stabilization of HSV-1 prefusion gB and binding of prefusion-specific antibodies. **(A)** SDS-PAGE of HSV-1 gB constructs under reducing and non-reducing conditions. The stabilizing interprotomeric disulfide bond shifts the band to a molecular weight consistent with that of a trimer. **(B)** Size-exclusion chromatography (SEC) trace of prefusion-stabilized HSV-1 gB and SDS-PAGE of pooled SEC fractions under reducing and non-reducing conditions. **(C)** Negative-stain electron microscopy 2D-class averages and 3D-reconstruction volumes of HSV-1 prefusion-stabilized gB complexed with Fab 3-6. **(D)** BLI sensorgrams for the association and dissociation kinetics of prefusion-specific and prefusion-preferring Fabs binding to immobilized HSV-1 prefusion-stabilized gB on Ni^2+^-NTA biosensors. Experimental data (black) were globally fit to a 1:1 binding model, with the fitted curves shown in red. The calculated values for the *K_d_* , *k_a_*, and *k_d_* are listed above each curve.

**Fig. S7.**
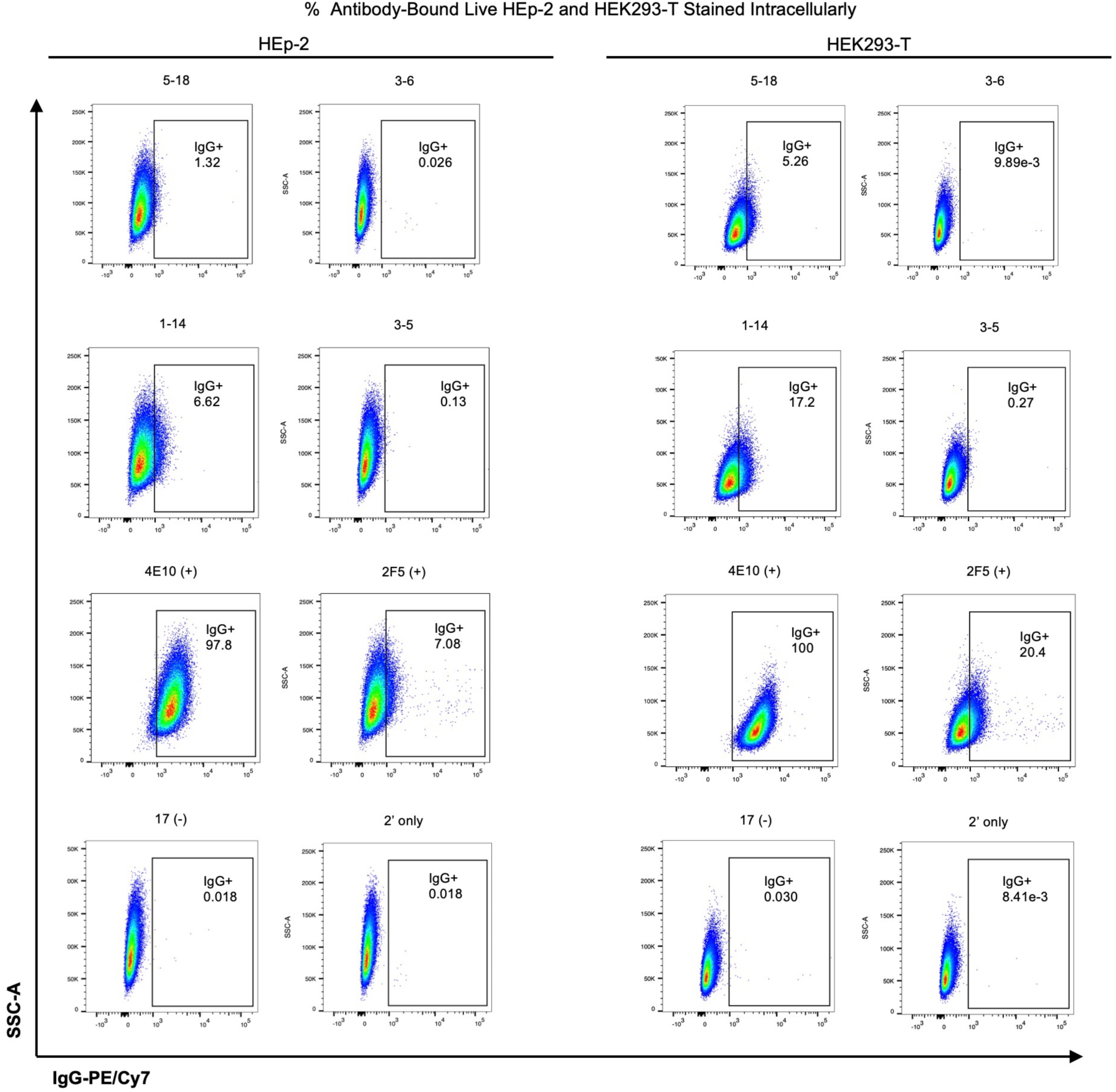
Flow cytometry plots of intracellular self-antigen recognition by prefusion-specific mAbs. Representative flow cytometry plots for intracellular staining of HEp-2 cells and HEK293-T cells with 25 µg/mL positive control mAbs 4E10 and 2F5, negative control 17B, or experimental mAbs 5-18, 3-6, 1-14, and 3-5. Anti-human IgG Fc-PE/Cy7 was used to detect experimental and control recognition of cellular autoantigens.

**Fig. S8.**
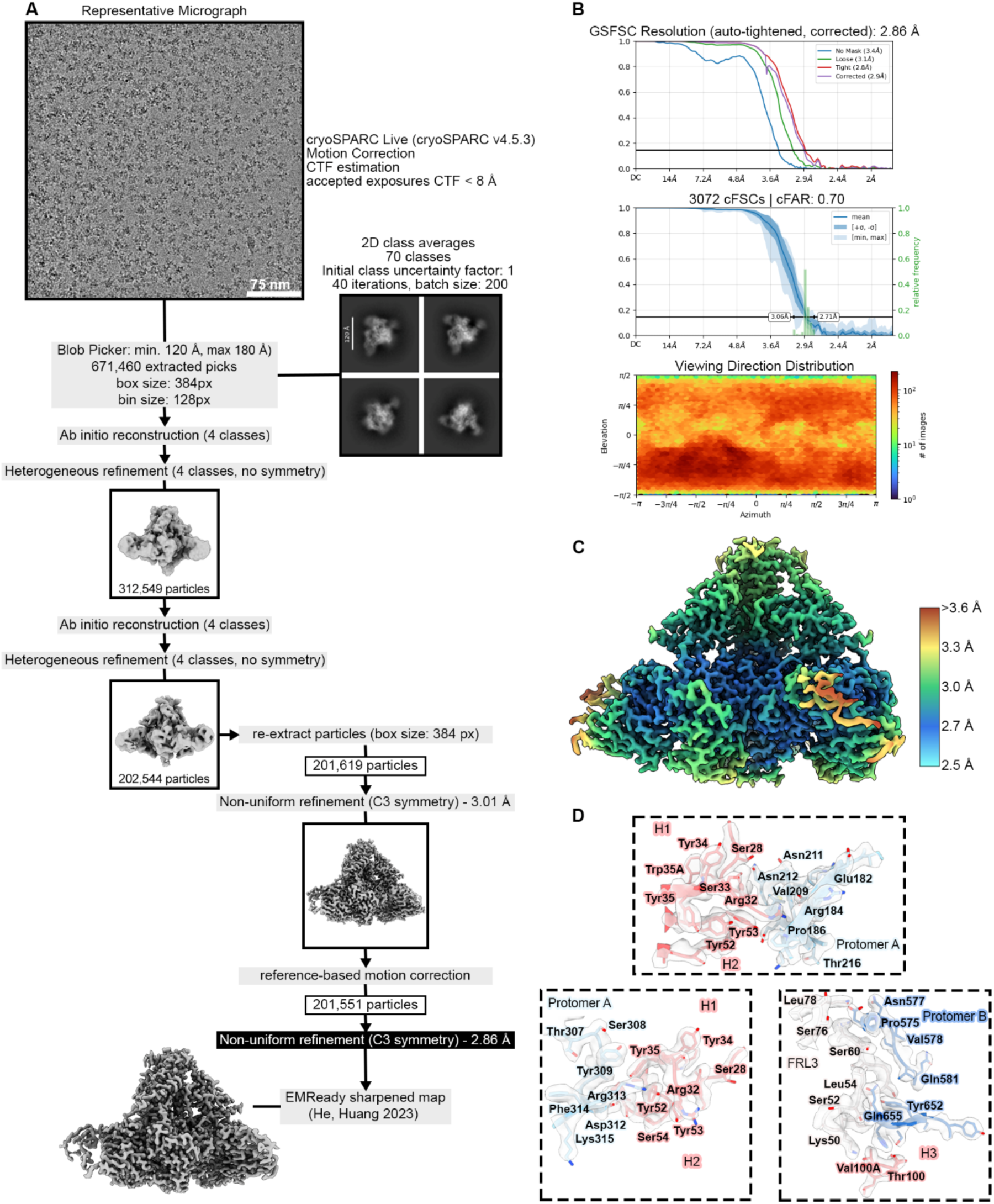
Cryo-EM data processing workflow and structure validation for prefusion HSV-2 gB bound to Fab 5-18.

**Fig. S9.**
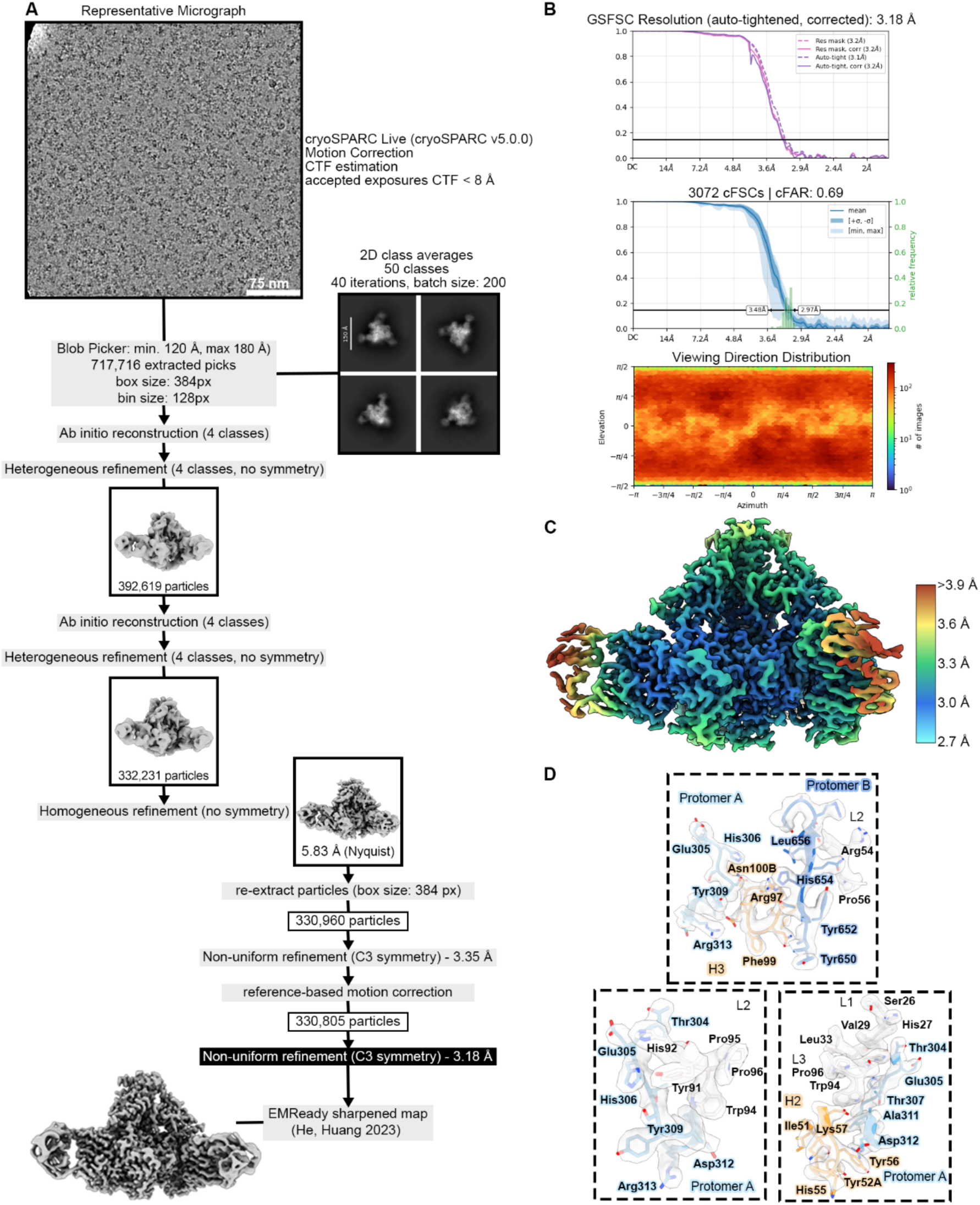
Cryo-EM data processing workflow and structure validation for prefusion HSV-2 gB bound to Fab 1-14.

**Fig. S10.**
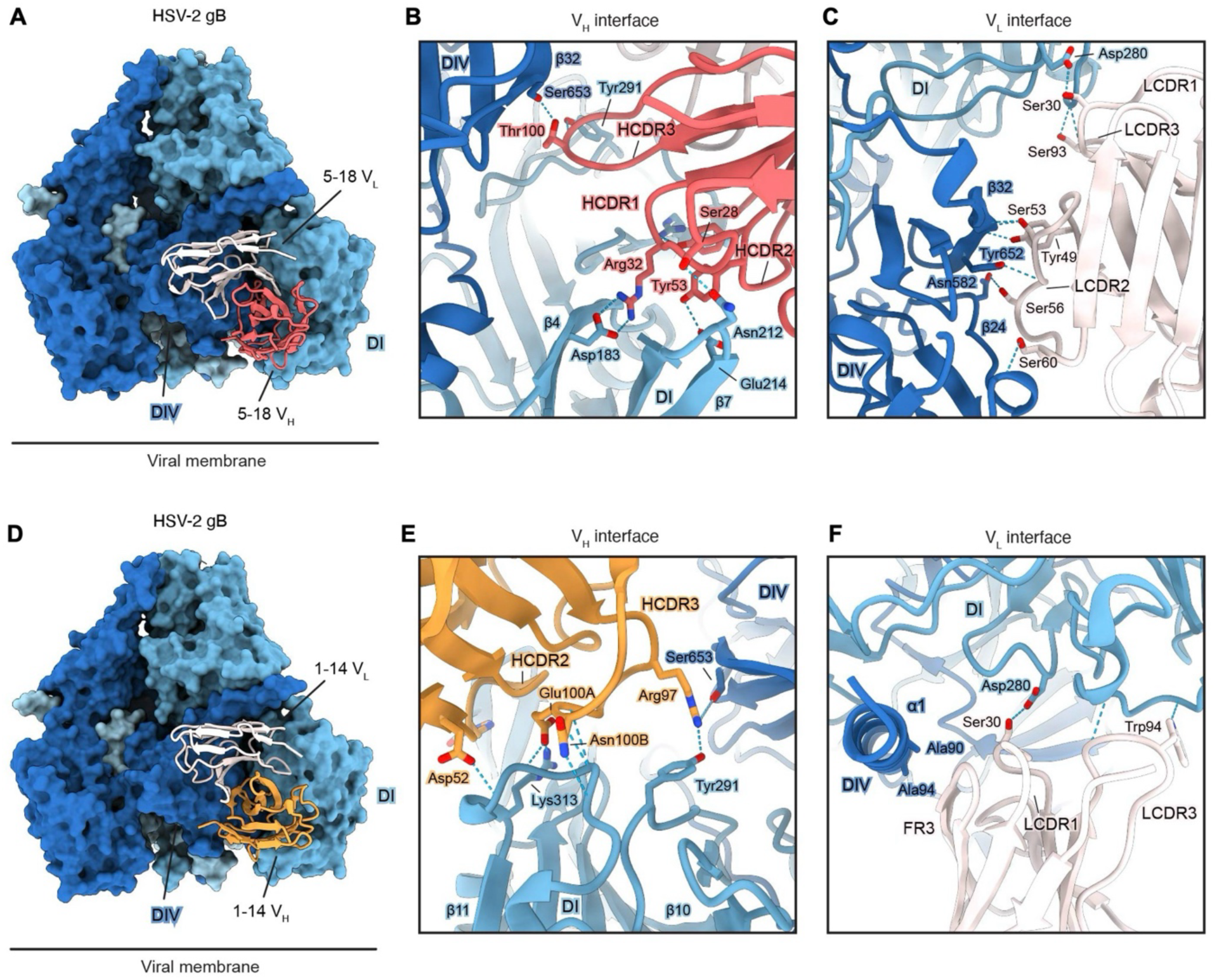
Quaternary interfaces of prefusion-specific Fabs 5-18 and 1-14. (A), Side view of prefusion HSV-2 gB with one bound Fab 5-18. A horizontal black line indicates the approximate position of the viral membrane. **(B)** Magnified view of the 5-18 VH interface spanning Domain (D) I and DIV, depicted as ribbons. HCDR1 and HCDR2 contact DI through multiple hydrogen bonds and salt bridges, with Arg32HCDR1 extending to engage the Asp183 sidechain in β4. HCDR3 spans DI and DIV, with Thr100HCDR3 forming hydrogen bonds to Tyr291 in DI and Ser653 in DIV. **(C)** Magnified view of the 5-18 VL interface spanning DI and DIV, depicted as ribbons. LCDR1 and LCDR3 interact with DI, whereas LCDR2 engages β24 and β32 in DIV, forming a continuous hydrogen bond network. **(D)** Side view of prefusion HSV-2 gB with one bound Fab 1-14. **(E)** Magnified view of the 1-14 VH interface spanning DI and IV, depicted as ribbons. HCDR2 and HCDR3 mediate most contacts with the β10–β11 loop in DI. HCDR3 crosslinks DI and DIV, with Arg97HCDR3 forming a hydrogen bond with the sidechain of Tyr291 in DI and Ser653 in DIV. **(F)** Magnified view of the 1-14 VL interface spanning DI and DIV, depicted as ribbons. LCDR1 and LCDR3 contact DI, whereas LCDR2 and FR3 make limited interactions with DIV, including the N-terminal α1 helix.

**Fig. S11.**
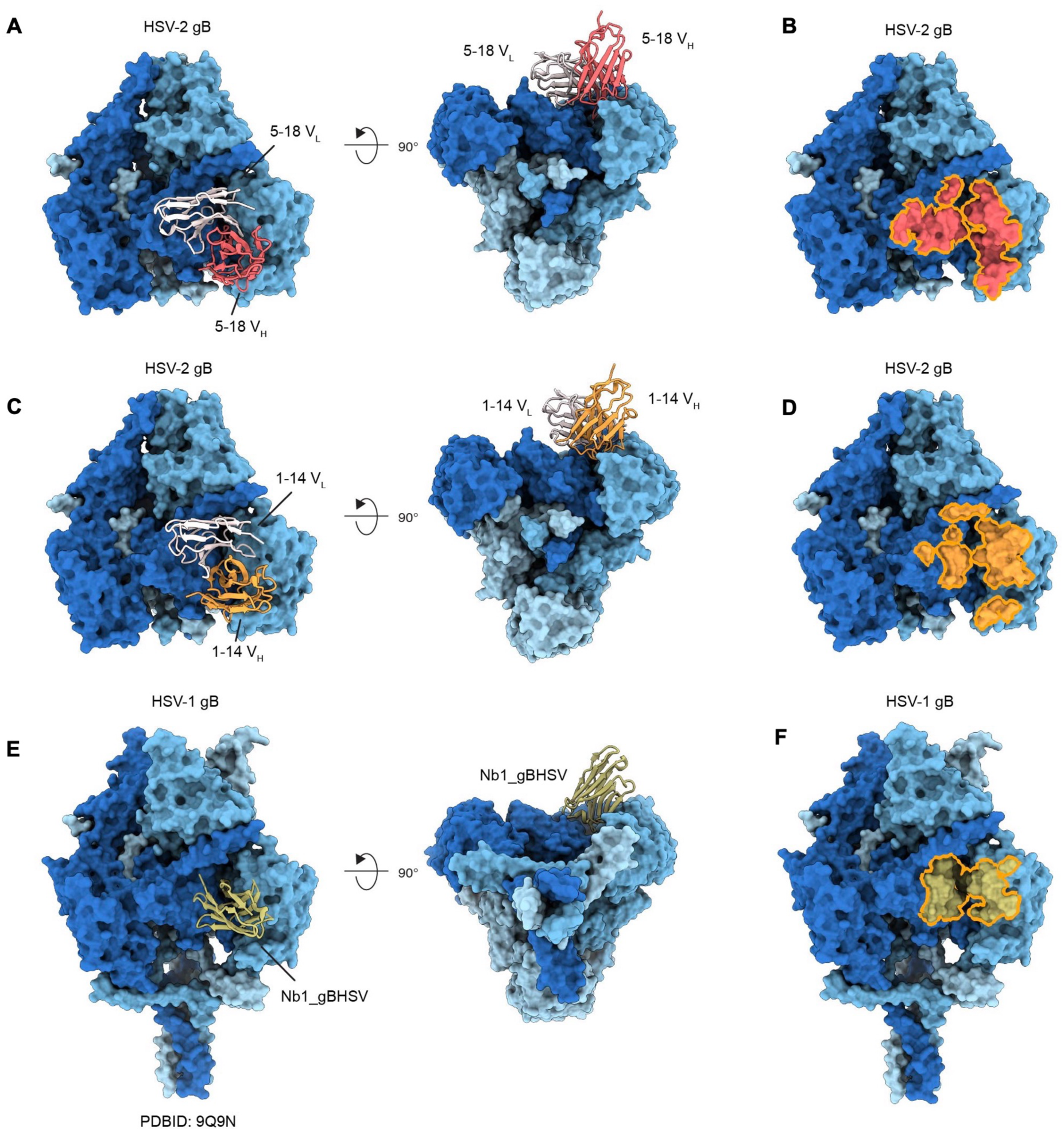
Structural comparison of prefusion-specific Fabs 5-18, 1-14, and nanobody Nb1_gBHSV. **(A)** Side (*left*) and bottom (*right*) views of prefusion HSV-2 gB bound to Fab 5-18. **(B)** Side view showing the Fab 5-18 epitope footprint outlined in orange. **(C)** Side (*left*) and bottom (*right*) views of prefusion HSV-2 gB bound to Fab 1-14. **(D)** Side view showing the Fab 1-14 epitope footprint. **(E)** Side (*left*) and bottom (*right*) views of prefusion HSV-1 gB bound to nanobody Nb1_gBHSV (PDBID: 9Q9N). **(F)** Side view showing the Nb1_gBHSV epitope footprint.

**Fig. S12.**
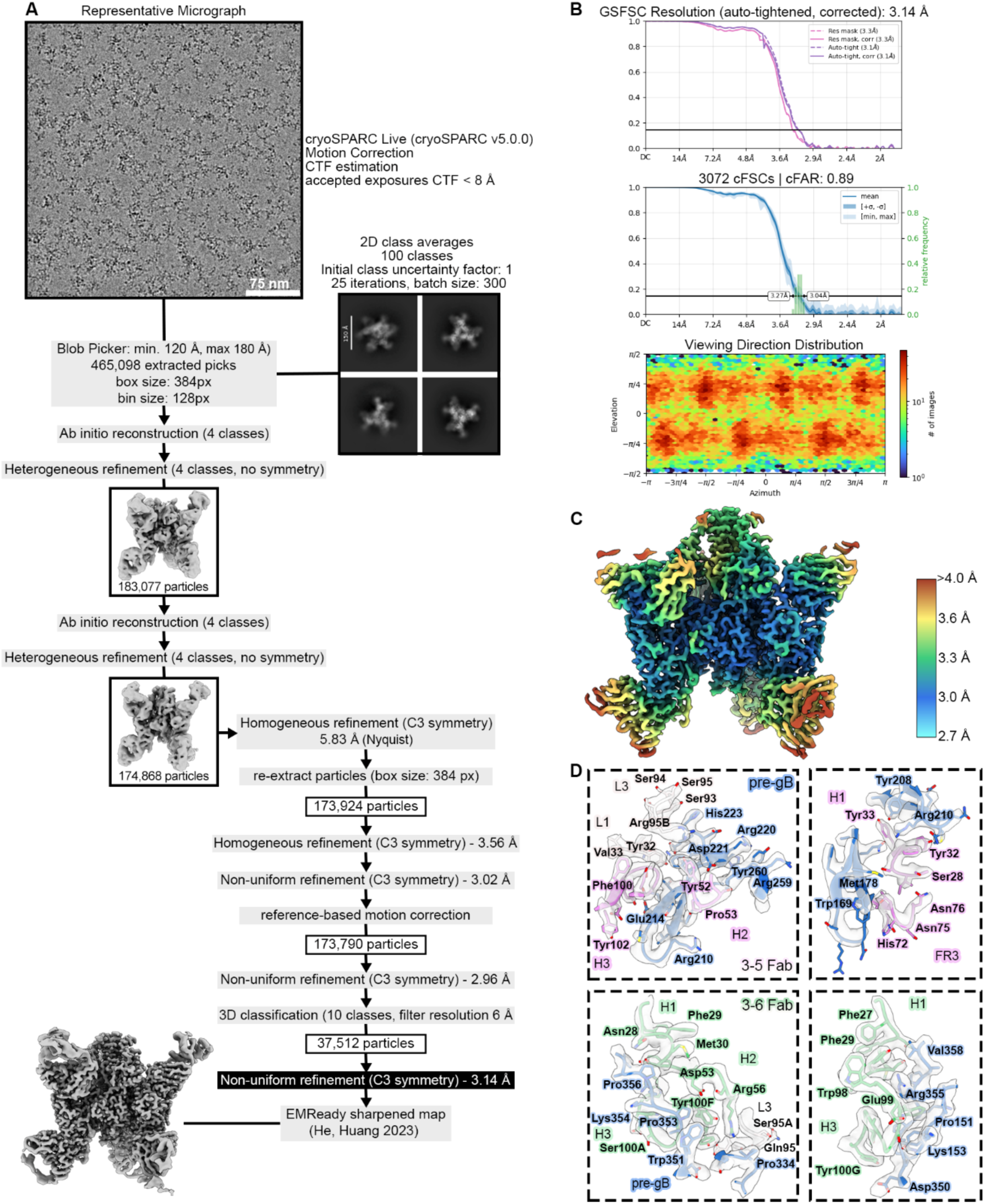
Cryo-EM data processing workflow and structure validation for prefusion HSV-2 gB bound to Fabs 3-6 and 3-5.

**Fig. S13.**
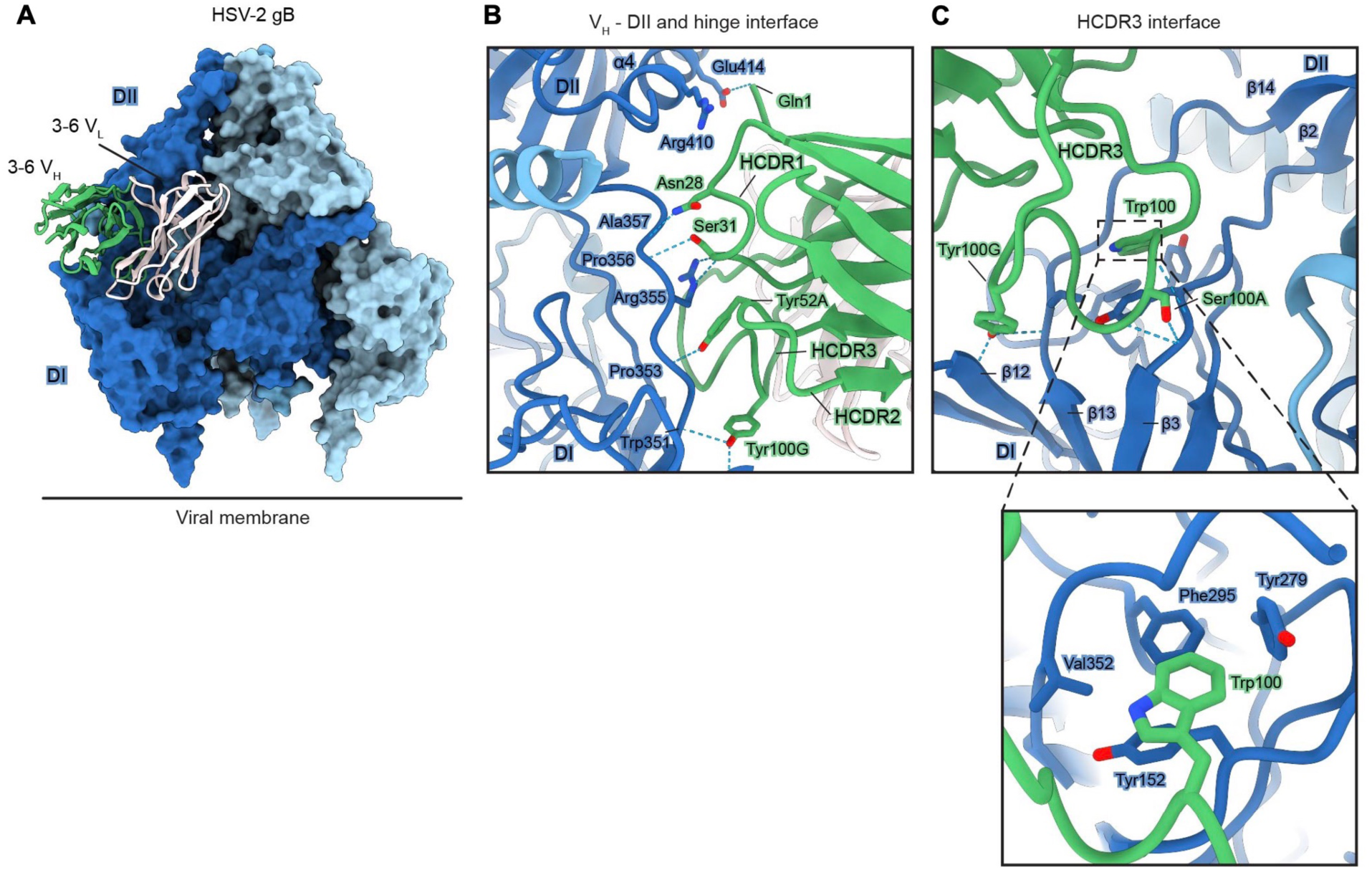
Interface of prefusion-specific Fab 3-6 with prefusion HSV-2 gB. **(A)** Side view of prefusion HSV-2 gB with one bound Fab 3-6. The 3-6 VH is colored green, the 3-6 VL is colored white, and each protomer of prefusion gB is colored in a different shade of blue. A horizontal black line indicates the approximate position of the viral membrane. **(B)** Magnified view of the 3-6 VH interface with the domain (D)II and hinge region, depicted as ribbons. FR1 and HCDR1 of the heavy chain interact with the α4 helix of DII. HCDR1, HCDR2, and HCDR3 form hydrogen bonds with mainchain atoms in hinge residues 351–358 and the sidechain of Arg355. **(C)** Magnified view of HCDR3 interactions with DI, depicted as ribbons. HCDR3 contacts β2–β3 loop, β12, and the β13–β14 loop (hinge). The inset shows Trp100HCDR3 inserted into a hydrophobic pocket formed by Tyr152, Tyr279, Phe295, and Val352.

**Fig. S14.**
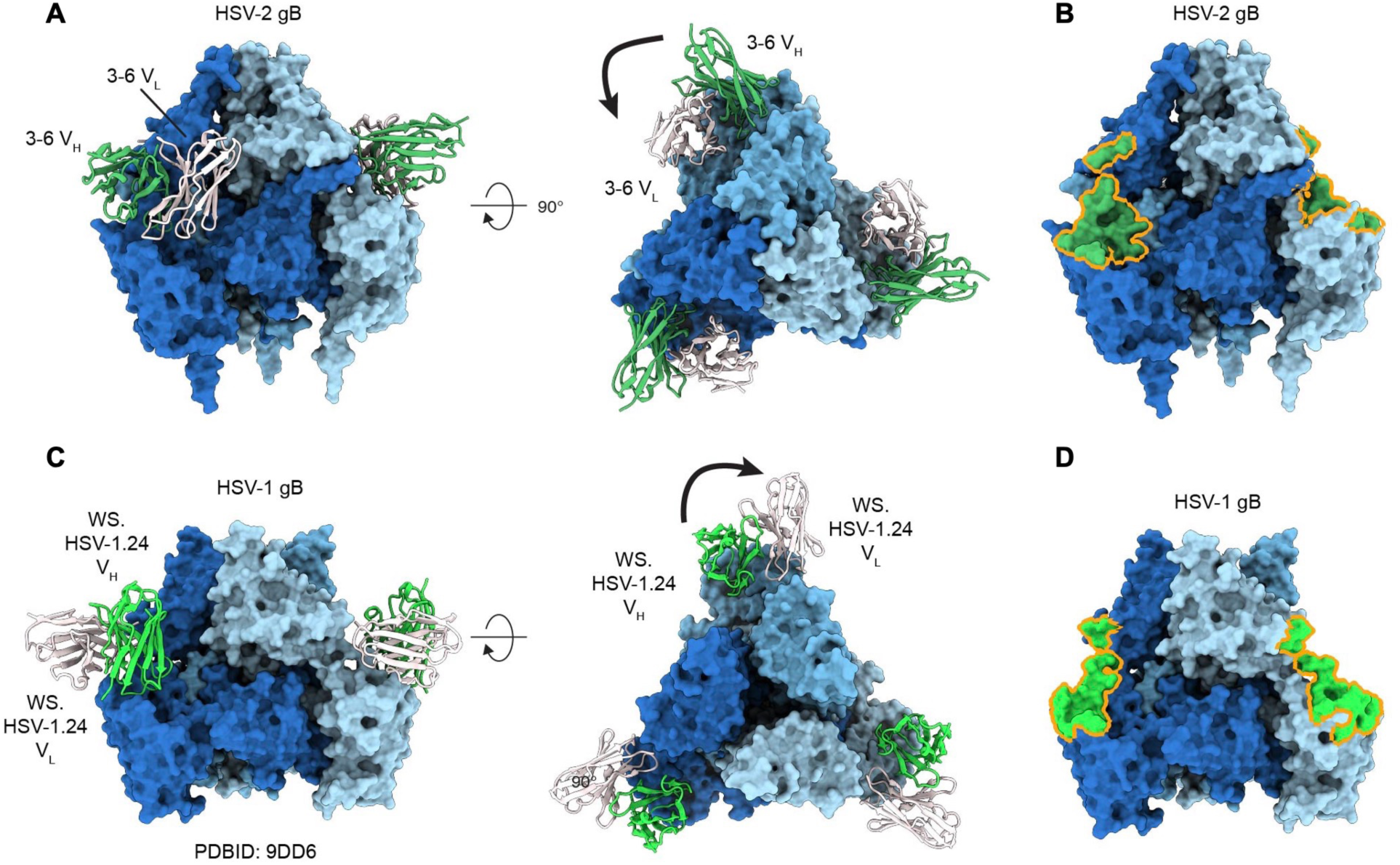
Structural comparison of prefusion-specific Fab 3-6 and murine mAb WS.HSV-1.24. **(A)** Side (left) and bottom (right) views of prefusion HSV-2 gB with three bound Fab 3-6. The 3-6 VH is colored green, the VL is colored white, and gB protomers are colored shades of blue. **(B)** Side view showing the epitope footprint of Fab 3-6 on prefusion HSV-2 gB. Footprints are outlined in orange. **(C)** Side (left) and bottom (right) views of prefusion HSV-1 gB with three bound Fab WS.HSV-1.24 (PDB ID: 9DD6). The WS.HSV-1.24 VH is colored light green, the VL is colored white, and gB protomers are colored shades of blue. **(D)** Side view showing the epitope footprint of Fab WS.HSV-1.24 on prefusion HSV-1 gB.

**Fig. S15.**
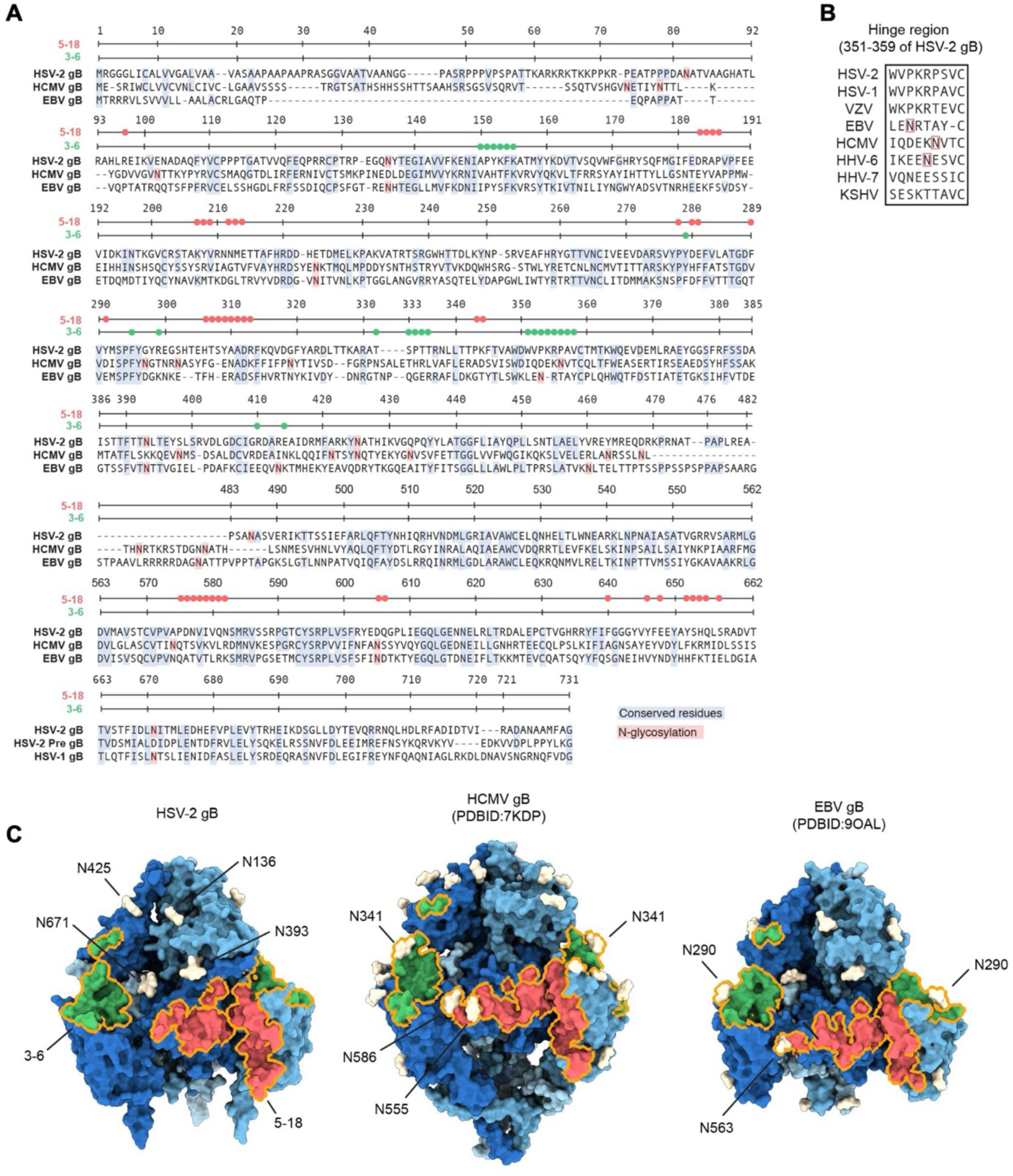
Conservation and glycan shielding of HSV-2 prefusion-specific epitopes across human herpesvirus gBs. **(A)** Sequence alignment of HSV-2 gB residues 1–731 (strain HG52), HCMV gB residues 1–711 (strain Towne), and EBV gB residues 1–692 (strain M81). Residues in the prefusion-specific epitopes of mAbs 5-18 (red) and 3-6 (green) are indicated above the alignments by colored circles. Conserved residues among HSV-2, HCMV, and EBV gB are highlighted in blue, and *N*-linked glycosylation sites are shown in red. **(B)** Sequence alignment of the domain (D)I–DII hinge-region residues from human herpesvirus gBs, with *N*-linked glycosylation sites in the hinge regions indicated by red boxes. **(C)** Structural comparison of the glycan shield near prefusion-specific epitopes in HSV-2, HCMV, and EBV gB. HSV-2 gB glycans are modeled based on the HSV-2 gB–Fab 3-5 and 3-6 complex structure. HCMV gB (PDB ID: 7KDP) and EBV gB (PDB ID: 9OAL) *N*-linked glycans are modeled using the GlycoShape server. The Fab 5-18 footprint is shown in red, the Fab 3-6 footprint in green, and both footprints are outlined in orange. Glycans are colored white. All built glycans are labeled for HSV-2, while only shielding glycans are labeled for HCMV and EBV gB

**Table S1.** Summary table of flow cytometry assay for intracellular and extracellular self-antigen recognition by prefusion-specific mAbs. Shown is the percent of antibody-bound HEp-2 and HEK293T live cell lines stained with mAbs intracellularly and extracellularly and gated for IgG-positive cells. Positive cutoff for autoreactivity set at 5% (bold).

| mAb | HEp-2 |  | HEK293-T |  |
| --- | --- | --- | --- | --- |
|  | Intracellular (%) | Extracellular (%) | Intracellular (%) | Extracellular (%) |
| 3-5 | 0.1 | 0.0 | 0.3 | 0.0 |
| 3-6 | 0.0 | 0.0 | 0.0 | 0.0 |
| 1-14 | **6.6** | 0.0 | **17.2** | 0.0 |
| 5-18 | 1.3 | 0.1 | **5.3** | 0.1 |
| 4E10 (+) | **97.8** | 0.0 | **100.0** | 0.0 |
| 2F5 (+) | **7.1** | 0.4 | **20.4** | 0.4 |
| 17B (-) | 0.0 | 0.0 | 0.0 | 0.0 |
| VRC01 (-) | **22.1** | 0.0 | **32.9** | 0.0 |
| 2' only | 0.0 | 0.0 | 0.0 | 0.0 |

**Table S2.**
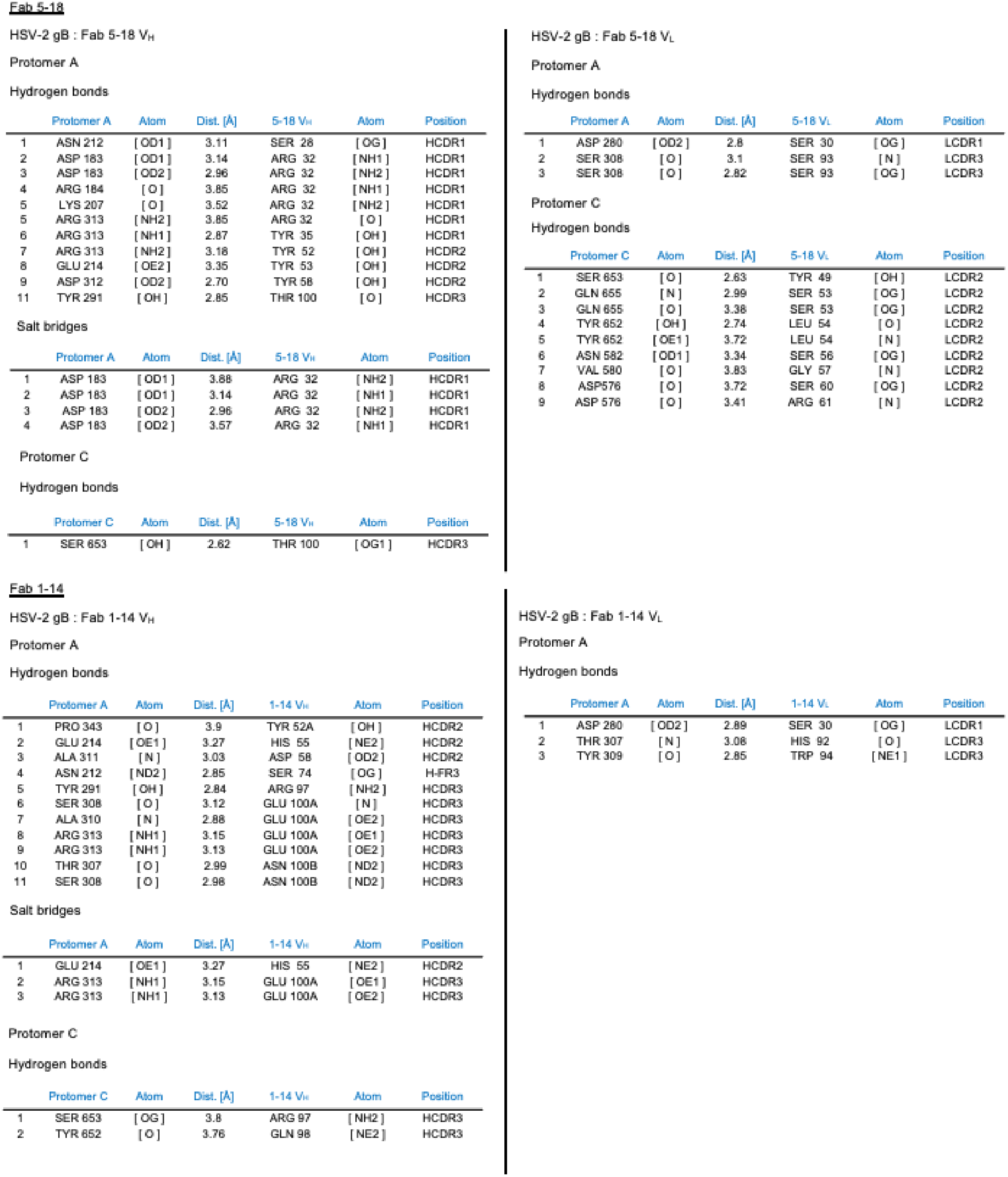
Interface analysis of mAbs 5-18 and 1-14 with prefusion stabilized HSV-2 gB. (PDBePISA)

|  | Protomer A | Atom | Dist. [Å] | 5-18 V<sub>H</sub> | Atom | Position |
| --- | --- | --- | --- | --- | --- | --- |
| 1 | ASN 212 | [OD1] | 3.11 | SER 28 | [OG] | HCDR1 |
| 2 | ASP 183 | [OD1] | 3.14 | ARG 32 | [NH1] | HCDR1 |
| 3 | ASP 183 | [OD2] | 2.96 | ARG 32 | [NH2] | HCDR1 |
| 4 | ARG 184 | [O] | 3.85 | ARG 32 | [NH1] | HCDR1 |
| 5 | LYS 207 | [O] | 3.52 | ARG 32 | [NH2] | HCDR1 |
| 5 | ARG 313 | [NH2] | 3.85 | ARG 32 | [O] | HCDR1 |
| 6 | ARG 313 | [NH1] | 2.87 | TYR 35 | [OH] | HCDR1 |
| 7 | ARG 313 | [NH2] | 3.18 | TYR 52 | [OH] | HCDR2 |
| 8 | GLU 214 | [OE2] | 3.35 | TYR 53 | [OH] | HCDR2 |
| 9 | ASP 312 | [OD2] | 2.70 | TYR 58 | [OH] | HCDR2 |
| 11 | TYR 291 | [OH] | 2.85 | THR 100 | [O] | HCDR3 |

Salt bridges
|  | Protomer A | Atom | Dist. [Å] | 5-18 V<sub>H</sub> | Atom | Position |
| --- | --- | --- | --- | --- | --- | --- |
| 1 | ASP 183 | [OD1] | 3.88 | ARG 32 | [NH2] | HCDR1 |
| 2 | ASP 183 | [OD1] | 3.14 | ARG 32 | [NH1] | HCDR1 |
| 3 | ASP 183 | [OD2] | 2.96 | ARG 32 | [NH2] | HCDR1 |
| 4 | ASP 183 | [OD2] | 3.57 | ARG 32 | [NH1] | HCDR1 |

Protomer C Hydrogen bonds
|  | Protomer C | Atom | Dist. [Å] | 5-18 V<sub>H</sub> | Atom | Position |
| --- | --- | --- | --- | --- | --- | --- |
| 1 | SER 653 | [OH] | 2.62 | THR 100 | [OG1] | HCDR3 |

|  | Protomer A | Atom | Dist. [Å] | 1-14 V<sub>H</sub> | Atom | Position |
| --- | --- | --- | --- | --- | --- | --- |
| 1 | PRO 343 | [O] | 3.9 | TYR 52A | [OH] | HCDR2 |
| 2 | GLU 214 | [OE1] | 3.27 | HIS 55 | [NE2] | HCDR2 |
| 3 | ALA 311 | [N] | 3.03 | ASP 58 | [OD2] | HCDR2 |
| 4 | ASN 212 | [ND2] | 2.85 | SER 74 | [OG] | H-FR3 |
| 5 | TYR 291 | [OH] | 2.84 | ARG 97 | [NH2] | HCDR3 |
| 6 | SER 308 | [O] | 3.12 | GLU 100A | [N] | HCDR3 |
| 7 | ALA 310 | [N] | 2.88 | GLU 100A | [OE2] | HCDR3 |
| 8 | ARG 313 | [NH1] | 3.15 | GLU 100A | [OE1] | HCDR3 |
| 9 | ARG 313 | [NH1] | 3.13 | GLU 100A | [OE2] | HCDR3 |
| 10 | THR 307 | [O] | 2.99 | ASN 100B | [ND2] | HCDR3 |
| 11 | SER 308 | [O] | 2.98 | ASN 100B | [ND2] | HCDR3 |

Salt bridges
|  | Protomer A | Atom | Dist. [Å] | 1-14 V<sub>H</sub> | Atom | Position |
| --- | --- | --- | --- | --- | --- | --- |
| 1 | GLU 214 | [OE1] | 3.27 | HIS 55 | [NE2] | HCDR2 |
| 2 | ARG 313 | [NH1] | 3.15 | GLU 100A | [OE1] | HCDR3 |
| 3 | ARG 313 | [NH1] | 3.13 | GLU 100A | [OE2] | HCDR3 |

Protomer C Hydrogen bonds
|  | Protomer C | Atom | Dist. [Å] | 1-14 V<sub>H</sub> | Atom | Position |
| --- | --- | --- | --- | --- | --- | --- |
| 1 | SER 653 | [OG] | 3.8 | ARG 97 | [NH2] | HCDR3 |
| 2 | TYR 652 | [O] | 3.76 | GLN 98 | [NE2] | HCDR3 |

|  | Protomer A | Atom | Dist. [Å] | 5-18 V<sub>L</sub> | Atom | Position |
| --- | --- | --- | --- | --- | --- | --- |
| 1 | ASP 280 | [OD2] | 2.8 | SER 30 | [OG] | LCDR1 |
| 2 | SER 308 | [O] | 3.1 | SER 93 | [N] | LCDR3 |
| 3 | SER 308 | [O] | 2.82 | SER 93 | [OG] | LCDR3 |

Protomer C Hydrogen bonds
|  | Protomer C | Atom | Dist. [Å] | 5-18 V<sub>L</sub> | Atom | Position |
| --- | --- | --- | --- | --- | --- | --- |
| 1 | SER 653 | [O] | 2.63 | TYR 49 | [OH] | LCDR2 |
| 2 | GLN 655 | [N] | 2.99 | SER 53 | [OG] | LCDR2 |
| 3 | GLN 655 | [O] | 3.38 | SER 53 | [OG] | LCDR2 |
| 4 | TYR 652 | [OH] | 2.74 | LEU 54 | [O] | LCDR2 |
| 5 | TYR 652 | [OE1] | 3.72 | LEU 54 | [N] | LCDR2 |
| 6 | ASN 582 | [OD1] | 3.34 | SER 56 | [OG] | LCDR2 |
| 7 | VAL 580 | [O] | 3.83 | GLY 57 | [N] | LCDR2 |
| 8 | ASP576 | [O] | 3.72 | SER 60 | [OG] | LCDR2 |
| 9 | ASP 576 | [O] | 3.41 | ARG 61 | [N] | LCDR2 |

|  | Protomer A | Atom | Dist. [Å] | 1-14 V<sub>L</sub> | Atom | Position |
| --- | --- | --- | --- | --- | --- | --- |
| 1 | ASP 280 | [OD2] | 2.89 | SER 30 | [OG] | LCDR1 |
| 2 | THR 307 | [N] | 3.08 | HIS 92 | [O] | LCDR3 |
| 3 | TYR 309 | [O] | 2.85 | TRP 94 | [NE1] | LCDR3 |

**Table S3.**
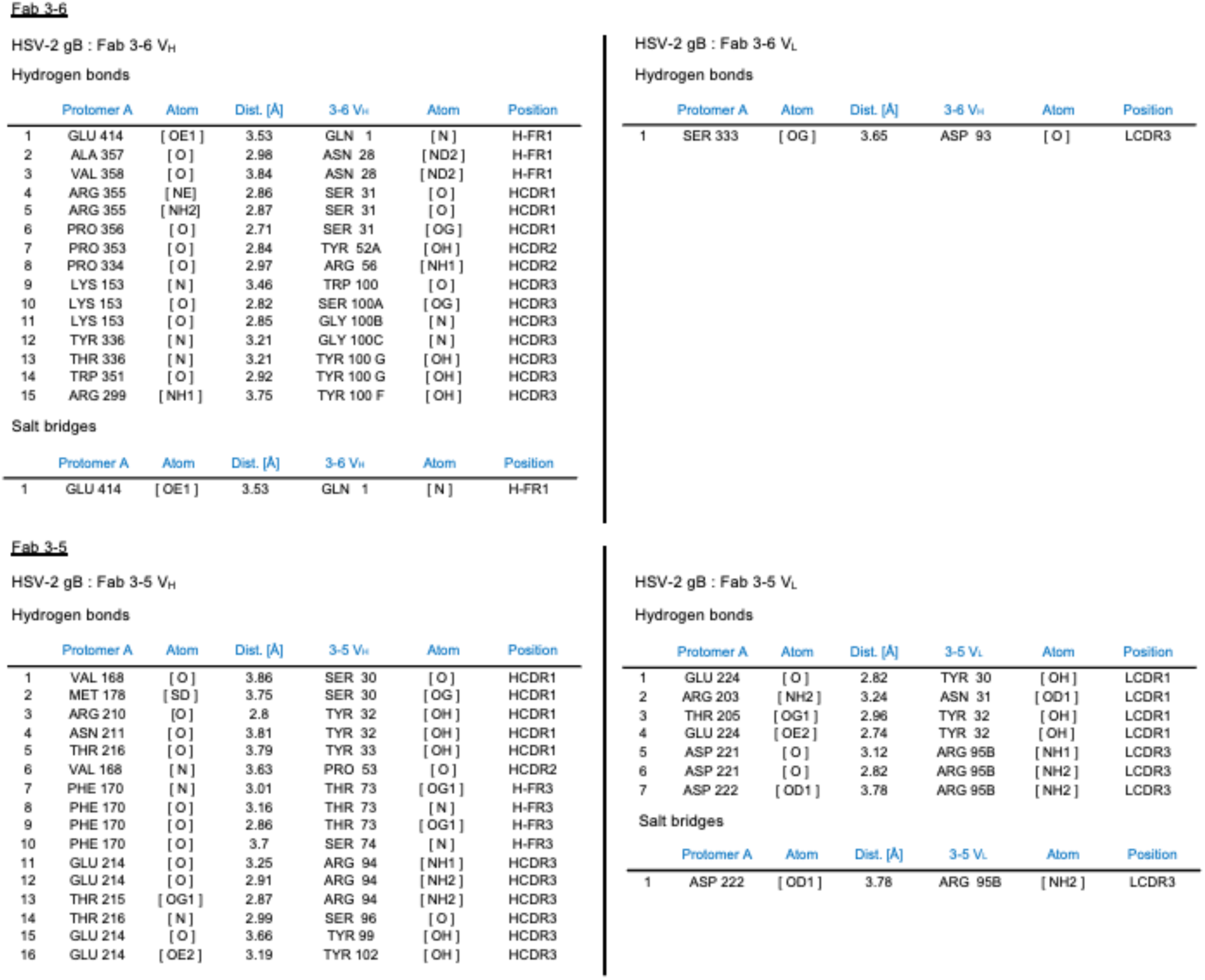
Interface analysis of mAbs 3-5 and 3-6 with prefusion-stabilized HSV-2 gB (PDBePISA).

|  | Protomer A | Atom | Dist. [Å] | 3-6 V<sub>H</sub> | Atom | Position |
| --- | --- | --- | --- | --- | --- | --- |
| 1 | GLU 414 | [OE1] | 3.53 | GLN 1 | [N] | H-FR1 |
| 2 | ALA 357 | [O] | 2.98 | ASN 28 | [ND2] | H-FR1 |
| 3 | VAL 358 | [O] | 3.84 | ASN 28 | [ND2] | H-FR1 |
| 4 | ARG 355 | [NE] | 2.86 | SER 31 | [O] | HCDR1 |
| 5 | ARG 355 | [NH2] | 2.87 | SER 31 | [O] | HCDR1 |
| 6 | PRO 356 | [O] | 2.71 | SER 31 | [OG] | HCDR1 |
| 7 | PRO 353 | [O] | 2.84 | TYR 52A | [OH] | HCDR2 |
| 8 | PRO 334 | [O] | 2.97 | ARG 56 | [NH1] | HCDR2 |
| 9 | LYS 153 | [N] | 3.46 | TRP 100 | [O] | HCDR3 |
| 10 | LYS 153 | [O] | 2.82 | SER 100A | [OG] | HCDR3 |
| 11 | LYS 153 | [O] | 2.85 | GLY 100B | [N] | HCDR3 |
| 12 | TYR 336 | [N] | 3.21 | GLY 100C | [N] | HCDR3 |
| 13 | THR 336 | [N] | 3.21 | TYR 100 G | [OH] | HCDR3 |
| 14 | TRP 351 | [O] | 2.92 | TYR 100 G | [OH] | HCDR3 |
| 15 | ARG 299 | [NH1] | 3.75 | TYR 100 F | [OH] | HCDR3 |

Salt bridges
|  | Protomer A | Atom | Dist. [Å] | 3-6 V<sub>H</sub> | Atom | Position |
| --- | --- | --- | --- | --- | --- | --- |
| 1 | GLU 414 | [OE1] | 3.53 | GLN 1 | [N] | H-FR1 |

|  | Protomer A | Atom | Dist. [Å] | 3-5 V<sub>H</sub> | Atom | Position |
| --- | --- | --- | --- | --- | --- | --- |
| 1 | VAL 168 | [O] | 3.86 | SER 30 | [O] | HCDR1 |
| 2 | MET 178 | [SD] | 3.75 | SER 30 | [OG] | HCDR1 |
| 3 | ARG 210 | [O] | 2.8 | TYR 32 | [OH] | HCDR1 |
| 4 | ASN 211 | [O] | 3.81 | TYR 32 | [OH] | HCDR1 |
| 5 | THR 216 | [O] | 3.79 | TYR 33 | [OH] | HCDR1 |
| 6 | VAL 168 | [N] | 3.63 | PRO 53 | [O] | HCDR2 |
| 7 | PHE 170 | [N] | 3.01 | THR 73 | [OG1] | H-FR3 |
| 8 | PHE 170 | [O] | 3.16 | THR 73 | [N] | H-FR3 |
| 9 | PHE 170 | [O] | 2.86 | THR 73 | [OG1] | H-FR3 |
| 10 | PHE 170 | [O] | 3.7 | SER 74 | [N] | H-FR3 |
| 11 | GLU 214 | [O] | 3.25 | ARG 94 | [NH1] | HCDR3 |
| 12 | GLU 214 | [O] | 2.91 | ARG 94 | [NH2] | HCDR3 |
| 13 | THR 215 | [OG1] | 2.87 | ARG 94 | [NH2] | HCDR3 |
| 14 | THR 216 | [N] | 2.99 | SER 96 | [O] | HCDR3 |
| 15 | GLU 214 | [O] | 3.66 | TYR 99 | [OH] | HCDR3 |
| 16 | GLU 214 | [OE2] | 3.19 | TYR 102 | [OH] | HCDR3 |

HSV-2 gB : Fab 3-6 V<sub>L</sub> Hydrogen bonds
|  | Protomer A | Atom | Dist. [Å] | 3-6 V<sub>L</sub> | Atom | Position |
| --- | --- | --- | --- | --- | --- | --- |
| 1 | SER 333 | [OG] | 3.65 | ASP 93 | [O] | LCDR3 |

HSV-2 gB : Fab 3-5 V<sub>L</sub> Hydrogen bonds
|  | Protomer A | Atom | Dist. [Å] | 3-5 V<sub>L</sub> | Atom | Position |
| --- | --- | --- | --- | --- | --- | --- |
| 1 | GLU 224 | [O] | 2.82 | TYR 30 | [OH] | LCDR1 |
| 2 | ARG 203 | [NH2] | 3.24 | ASN 31 | [OD1] | LCDR1 |
| 3 | THR 205 | [OG1] | 2.96 | TYR 32 | [OH] | LCDR1 |
| 4 | GLU 224 | [OE2] | 2.74 | TYR 32 | [OH] | LCDR1 |
| 5 | ASP 221 | [O] | 3.12 | ARG 95B | [NH1] | LCDR3 |
| 6 | ASP 221 | [O] | 2.82 | ARG 95B | [NH2] | LCDR3 |
| 7 | ASP 222 | [OD1] | 3.78 | ARG 95B | [NH2] | LCDR3 |

|  | Protomer A | Atom | Dist. [Å] | 3-5 V<sub>L</sub> | Atom | Position |
| --- | --- | --- | --- | --- | --- | --- |
| 1 | ASP 222 | [OD1] | 3.78 | ARG 95B | [NH2] | LCDR3 |

**Table S4.**
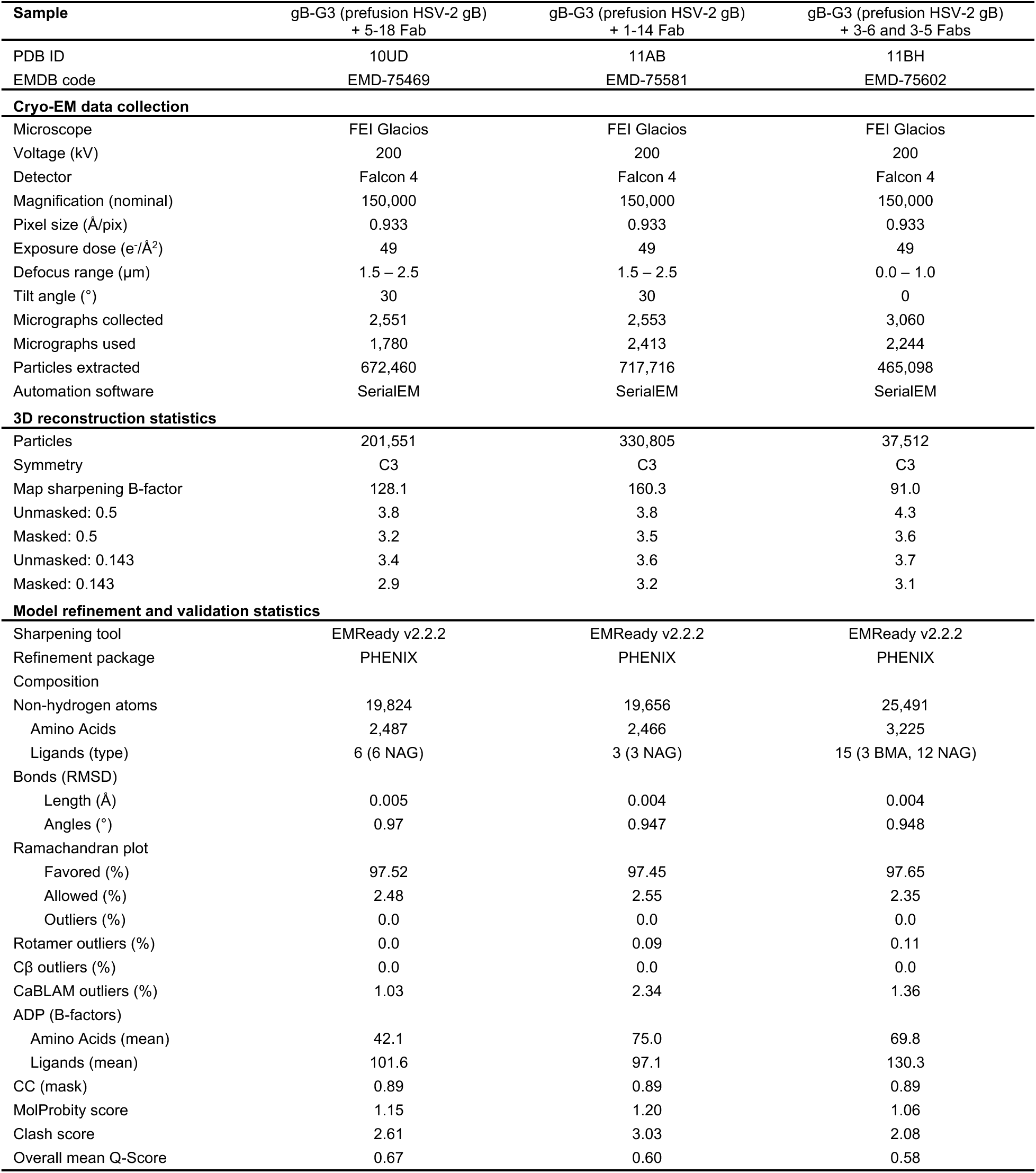
Cryo-EM Data collection and refinement statistics.

